# MultiOMICs landscape of SARS-CoV-2-induced host responses in human lung epithelial cells

**DOI:** 10.1101/2022.09.06.506768

**Authors:** Sneha M. Pinto, Yashwanth Subbannayya, Hera Kim, Lars Hagen, Maria W. Górna, Anni I. Nieminen, Magnar Bjørås, Terje Espevik, Denis Kainov, Richard K. Kandasamy

**Author notes:** Correspondence to: Richard K. Kandasamy, Sneha M. Pinto.

## Abstract

Despite the availability of vaccines and approved therapeutics, the COVID-19 pandemic continues to rise owing to the emergence of newer variants. Several multi-omics studies have made available extensive evidence on host-pathogen interactions and potential therapeutic targets. Nonetheless, an increased understanding of host signaling networks regulated by post-translational modifications and their ensuing effect on the biochemical and cellular dynamics is critical to expanding the current knowledge on the host response to SARS-CoV-2 infections. Here, employing unbiased global transcriptomics, proteomics, acetylomics, phosphoproteomics, and exometabolome analysis of a lung-derived human cell line, we show that SARS-CoV-2 Norway/Trondheim-S15 strain induces time-dependent alterations in the induction of type I IFN response, activation of DNA damage response, dysregulated Hippo signaling, among others. We provide evidence for the interplay of phosphorylation and acetylation dynamics on host proteins and its effect on the altered release of metabolites, especially organic acids and ketone bodies. Together, our findings serve as a resource of potential targets that can aid in designing novel host-directed therapeutic strategies.

## Introduction

The rapid emergence of the COVID-19 pandemic caused by severe acute respiratory syndrome coronavirus 2 (SARS-CoV-2) still shows no abatement ^1^. Despite the availability of several vaccines, the emergence of newer variants with enhanced transmissibility and pathogenesis is a major cause of concern. Current efforts to develop antiviral therapeutics, especially host-directed therapies, are vital to significantly reduce the impact of both the current and future coronavirus epidemics. Several classes of FDA-approved drugs and drugs currently in clinical trials have been repurposed, showing promising results in hospitalized patients with severe diseases ^2, 3, 4, 5^.

Host-directed therapeutics development largely depends on the available information as the viral genome continuously evolves to adapt to the host environment and evade pre or post-countermeasures. Several host factors that promote or restrict SARS-CoV-2 replication have been identified by genome-wide CRISPR knockout screens ^6, 7, 8, 9, 10, 11, 12, 13, 14^. Further, multi-OMICS studies have provided vital insights into understanding the viral profile, mechanisms of pathogenesis, and identifying host factors ^15, 16, 17, 18, 19, 20, 21, 22, 23^. Notably, the role of protein post-translational modifications (PTMs), are increasingly recognized as key mechanisms viruses employ to target signaling pathways regulating host immune responses^24, 25, 26^. Nevertheless, strain-specific differences and their effects on the host cellular machinery, especially in evading the immune response and facilitating further transmissibility and clinical manifestations, are only recently being explored ^27^.

Given the evolving number of variants, a multi-OMICS approach focusing on the dynamic interplay of PTMs is vital as it aids in therapeutics development and is helpful in pandemic preparedness. Most studies have focused on elucidating the phosphorylation dynamics and alterations in ubiquitin and glycosylation ^18, 20, 27, 28, 29, 30, 31^. Some are restricted to characterizing the glycosylation patterns of virus/membrane proteins ^32, 33, 34, 35^. Emerging evidence highlights protein lysine acetylation as a key regulatory mechanism originally thought to modulate epiproteome. Epigenetic responses are now increasingly identified as promising therapeutic targets in human viral infections ^36, 37, 38^. However, alterations in global host acetylation dynamics and the dynamic interplay with other PTMs in mediating viral pathogenesis in response to SARS-CoV-2 infection remain to be explored. Further, the effect of the altered signaling dynamics on the metabolite profile and, in turn, its impact on cellular interactions upon infection is yet to be determined. Increasing evidence shows that in addition to altering levels of intracellular metabolites, viral infections can result in the altered release of metabolites into the extracellular milieu ^39^. Quantitative measurements of exometabolites or metabolic footprinting enable the determination of metabolic state, serving as a reporter of the cellular physiology ^40, 41, 42^. Given the vital significance of cell-cell interactions and the control of its microenvironment, it is important to characterize the exometabolome changes upon SARS-CoV-2 infection.

Here we present a quantitative survey of the alterations in the global transcriptome, proteome, phosphorylation, lysine acetylation and exometabolite dynamics of SARS-CoV-2 Norway/Trondheim-S15 strain infected Calu-3 human lung epithelial cells. We map the altered changes to disrupted kinases and signaling pathways affecting cellular response to infection and cell survival. Overall, our findings demonstrate the control of host machinery by SARS-CoV-2 through PTM modulation.

## Results and discussion

### 1. Quantitative temporal viromics analysis of SARS-CoV-2 infection

Host responses to SARS-CoV-2 infection were investigated using a multipronged approach comprising transcriptomics, TMT-based quantitative temporal proteomics, phosphoproteomics, acetylomics, and targeted exometabolomics analysis. The analysis was performed on human lung epithelial Calu-3 cells infected with a clinical isolate of SARS-CoV-2 (hCoV-19/Norway/Trondheim-S15/2020 strain) at an MOI of 0.1 at five-time points (3, 6-, 12-, 24-, and 48 hours post-infection (hpi) (**Figure 1A**). We chose this MOI as a productive viral infection, and increased cytopathic effects were observed 72 hours post-infection in a previous study that used the same viral strain ^43^. Robust changes in transcript abundance and protein PTM dynamics were observed compared to protein abundance changes, and the findings are in concordance with previous reports ^27^. Across the datasets, 17,663 transcripts, 6,080 proteins, 24,013 phosphorylated peptides mapping to 5,487 phosphoproteins, 1,503 acetylated peptides mapping to 662 acetylated proteins, and 85 targeted metabolites were identified and quantified. A total of 3,457 transcripts, 216 proteins, 4,497 phosphopeptides mapping to 2,157 phosphoproteins, and 112 acetylated peptides mapping to 90 proteins, respectively, were found to be differentially regulated across all time points (fold change cutoff log2 >=1.5 or <=-1.5 for transcriptome and >=0.5 or <=-0.5 for other datasets) (**Figure 1B, Supplementary Tables S1-S4**).

**Figure 1.**
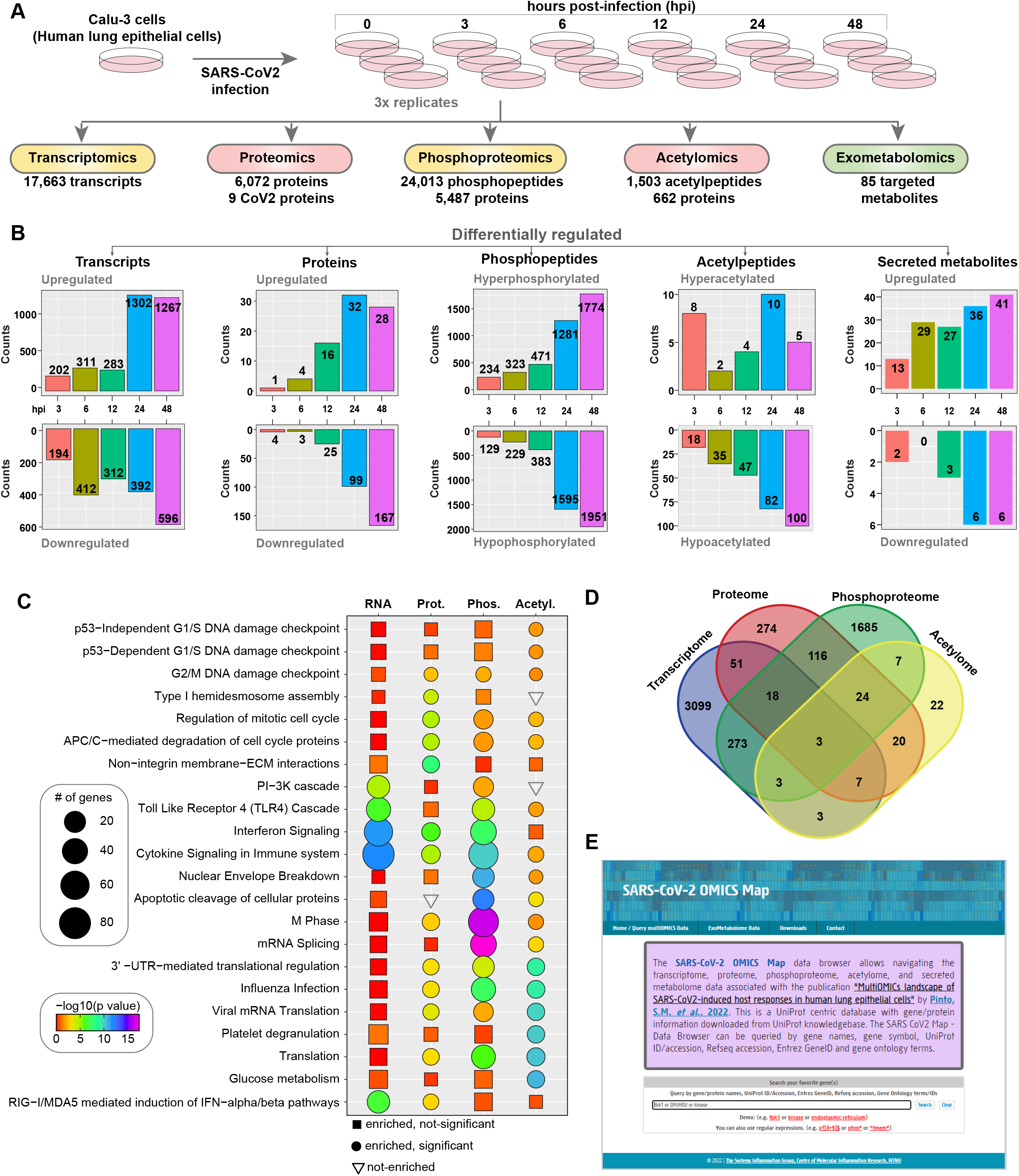
Multi-omics analysis of SARS-CoV-2 infected Calu-3 cells. **A.** Experimental workflow for the proteomic and phosphoproteomic analysis of Calu-3 cells infected with SARS-CoV-2/ Trondheim-S15/2020 strain. **B.** Result summary of differentially regulated transcripts, proteins, phosphopeptides, acetylpeptides, and secreted metabolites identified in Calu-3 cells infected with SARS-CoV-2. **C.** Heatmap showing enriched pathways for transcriptome, proteome, phosphoproteome, and acetylome data of SARS-CoV-2 Calu-3 cells. **D.** Overlap of genes across the different datasets from the current study. **E.** A screenshot of the SARS-CoV-2 OMICS Map containing data acquired in the current study.

Additionally, increased expression of viral genes at both the transcript and protein levels, as well as altered phosphorylation and acetylation levels in 7 viral proteins (acetylation observed only on nucleoprotein), was observed. Our data demonstrated a strong correlation between the time points and biological replicates post-data filtering and quality control analysis (**Figure S1A-D**). The extent of differential regulation of host and viral proteins was minimal in contrast to the marked hyperphosphorylation observed at earlier time points of infection. Contrarily, the acetylome dynamics revealed a profound downregulation as early as 6 hpi indicative of viral-induced PTM-specific regulatory mechanisms. Overall, enrichment of antiviral and innate immune signaling pathways and regulation of DNA damage response and cell cycle processes were observed across the different levels (**Figure 1C**). A minimal overlap across the differentially expressed transcript/protein/phosphoprotein/acetyl protein levels across all time points indicates discreet molecular-level regulation **(Figure 1D**). The multi-omics data is available as a queriable database “SARS-CoV-2 map” (http://www.sarsCoV-2map.org/index.html, **Figure 1E)**

### 2. Global proteome and transcriptome analysis reveal induction of innate immune response

Analysis of the global gene expression dynamics revealed a similar extent of gene regulation at early time points (3, 6, and 12 hpi), albeit with a slight increase in the number of downregulated genes at 12 hpi. Several genes were overexpressed at later time points of infection. Akin to transcriptome changes, host cellular proteins were significantly downregulated, especially 12 hpi (**Figure 2A, Supplementary Tables S1-S2)**. Our findings correlate with the shutoff of host mRNA translation activity and possibly cell cycle arrest, as observed in viral infections. In line with previous reports ^27, 29, 31, 44, 45^, significant enrichment of interferon and antiviral response processes at transcriptome and proteome levels was observed (**Figure S2A, Figure 1C**) despite a minimum overlap of differentially expressed proteins across the datasets derived from lung cell lines (**Figure S2B)**. Specifically, interferons-*IFNB1* (Type I IFN) and *IFNL1, −2, −3, −4* (Type III IFN) were induced as early as 12 hpi along with the overexpression of antiviral factors, and interferon-stimulated genes (ISGs) such as *OAS1-3, IFIT1-3, IFITM* at both transcriptomes and proteome levels (**Figure 2B-C**). Notably, CMPK2, an immunomodulatory ISG associated with antiviral responses ^46, 47, 48, 49^ in immune cells, and a kinase involved in mitochondrial DNA synthesis ^50^ and DNA repair ^51^, was upregulated at 24 and 48 hpi both at transcriptome and proteome levels. Using immunoblot analysis, we confirmed the induction of OAS1, ISG15, TRIM5α, RNase-L, MX1, and CMPK2. (**Figure 2D**). Comparing the ISG expression profile with that of the early lineage and Alpha variant ^27^ revealed a similar extent of expression in the Trondheim strain and Alpha variant compared to the early lineage viruses (IC19 and VIC), indicating strain-specific differences in inducing interferon response (**Figure 2E**).

**Figure 2.**
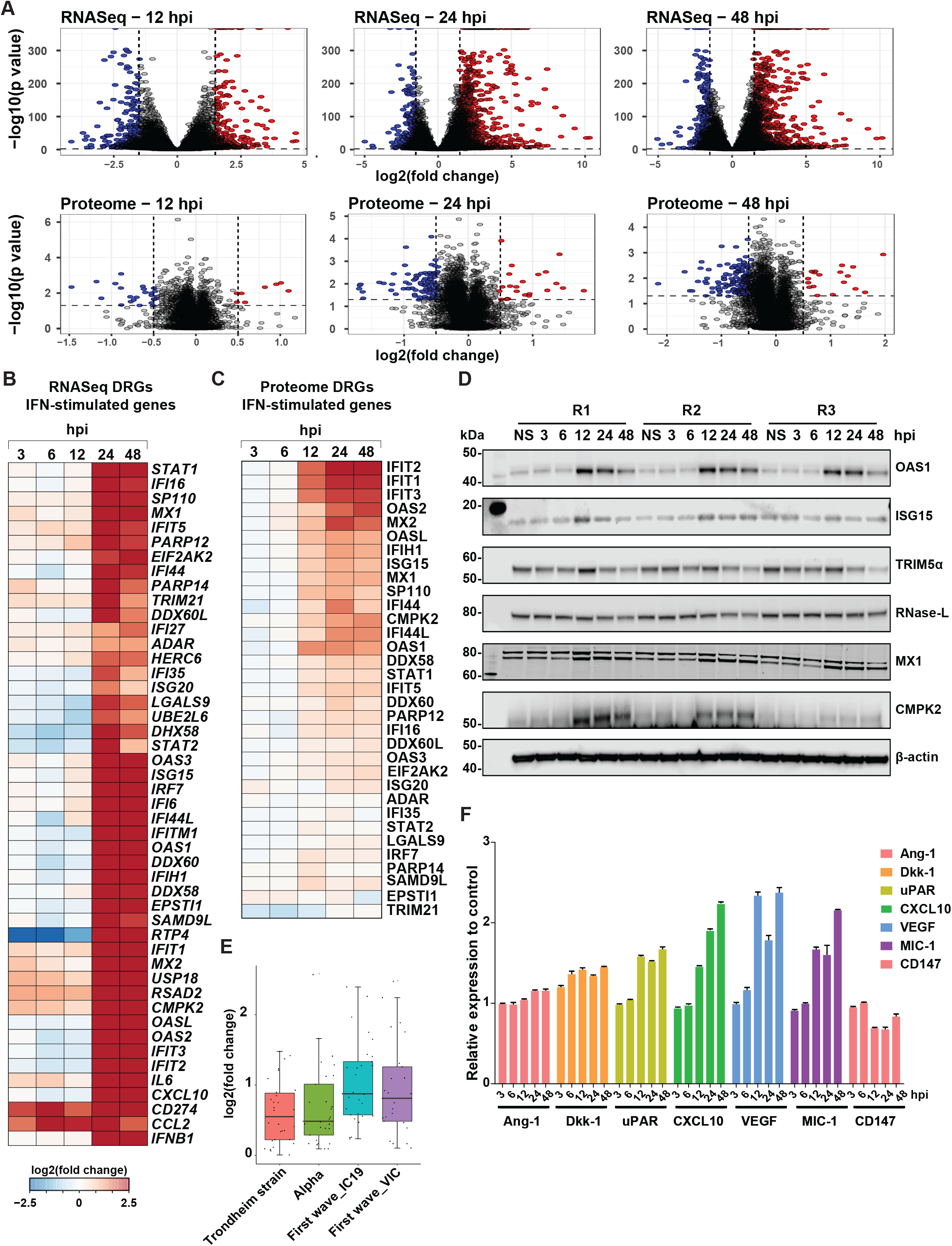
Proteomics and transcriptomic analysis of SARS-CoV-2 infected Calu-3 cells. **A.** Volcano plots displaying differential changes in RNA and protein expression of Calu-3 cells after infection with SARS-CoV-2/ Trondheim-S15/2020 strain at 12, 24, and 48 hpi. Red-filled circles indicate overexpressed genes/proteins, whereas blue-filled circles indicate downregulated genes/proteins. Heatmaps depicting differentially expressed IFN-stimulated genes from the **B.** RNA-Seq and **C.** Proteomics data of SARS-CoV-2-infected Calu-3 cells at 3, 6, 12, 24, and 48 hpi. **D.** Immunoblot analysis depicting time-dependent changes in the expression of interferon signaling proteins -OAS1, ISG15, TRIM5α, RNase-L, MX1, and CMPK2 in SARS-CoV-2-infected Calu-3 cells. **E.** A graph comparing transcript levels of interferon-stimulated genes (ISGs) between the Trondheim strain (S15), alpha, IC19, and VIC strains from Thorne *et al*. (2022) **F.** Cytokine array data showing differential levels of Ang-1, Dkk-1, uPAR, CXCL10, VEGF, MIC-1, and CD147 in Calu-3 cells post-infection.

While it is well known that Interferons activate Toll-like receptor (TLR) gene expression in macrophages during viral infections ^52^, including in SARS-CoV-2, expression of TLR genes in response to viral infections in epithelial cells is less known. Several members of the TLR family, including TLR1 (24 and 48 hpi), TLR3 (24 and 48 hpi), TLR4 (6 and 24 hpi), and TLR6 (24 and 48 hpi), were found to be upregulated in the Calu-3 transcriptome in response to infection. TLR2 has been reported to sense the SARS-CoV-2 envelope protein and thereby produce inflammatory cytokines, causing inflammation and damage in the lungs ^53^. Although we did not identify differential expression of TLR2, increased expression of TLR1 and TLR6, known to form heterodimers with TLR2 ^54^, was observed. While TLR3 activation has been reported in SARS-CoV-2-infected Calu-3 lung epithelial cells ^55^, there is indirect evidence that TLR4 may play a role in innate immune response in epithelial cells during SARS-CoV-2 infection ^56, 57^. Concerning the expression of the pro-inflammatory cytokine signature (*TNF, IL1B, IL6, CCL2, CXCL9, CXCL10, CXCL11*) previously shown to be upregulated in plasma and/or BAL of severe COVID-19 patients ^58, 59^, overexpression of *IL1B* was observed at early time points of infection (3-12 hpi), whereas the rest were upregulated at later time points (24-48 hpi) (**Figure S2C**). However, analysis of the collected supernatants indicated a time-dependent increase in CXCL10, VEGF, and GDF15 (MIC-1) levels, with no secretion of other pro-inflammatory cytokines. (**Figure 2F, Figure S2D**). Overall, our findings concord with previous studies indicating that Calu-3 cells respond to SARS-CoV-2 infection and induce an anti-viral response, albeit to a varying extent compared to other strains.

### 3. SARS-CoV-2 mediated modulation of PTM dynamics affects key host signaling pathways contributing to viral pathogenesis

To assess the impact of SARS-CoV-2 infection on host cellular signaling dynamics, serial enrichment analysis of phosphorylation and lysine acetylation was performed. Supervised clustering of the global Calu-3 phosphoproteome dynamics in response to SARS-CoV-2 infection revealed 8 distinct clusters corresponding to early (clusters 5 and 6), intermediate (cluster 6), late (hyperphosphorylated: clusters 1, 7, and 8; hypophosphorylated: cluster 2), and sustained responses (clusters 3 and 4) (**Figure 3A**). A minimal overlap was observed when comparing the hyperphosphorylated sites identified in the current study with other studies on the SARS-CoV-2 phosphoproteome ^27, 29, 31^ (**Figure S3A**). Given the differences in the cell types, strains and platforms, it is not surprising that a large majority of sites identified vary across studies. Nevertheless, pathway analysis revealed enrichment of cytokine signaling, activation of DNA damage and repair pathways, Hippo signaling, cell cycle regulation, RNA splicing, and regulation of translation and transcription across different clusters indicative of viral-mediated alterations of host cellular machinery (**Figure 3B, Supplementary Table S5**).

**Figure 3.**
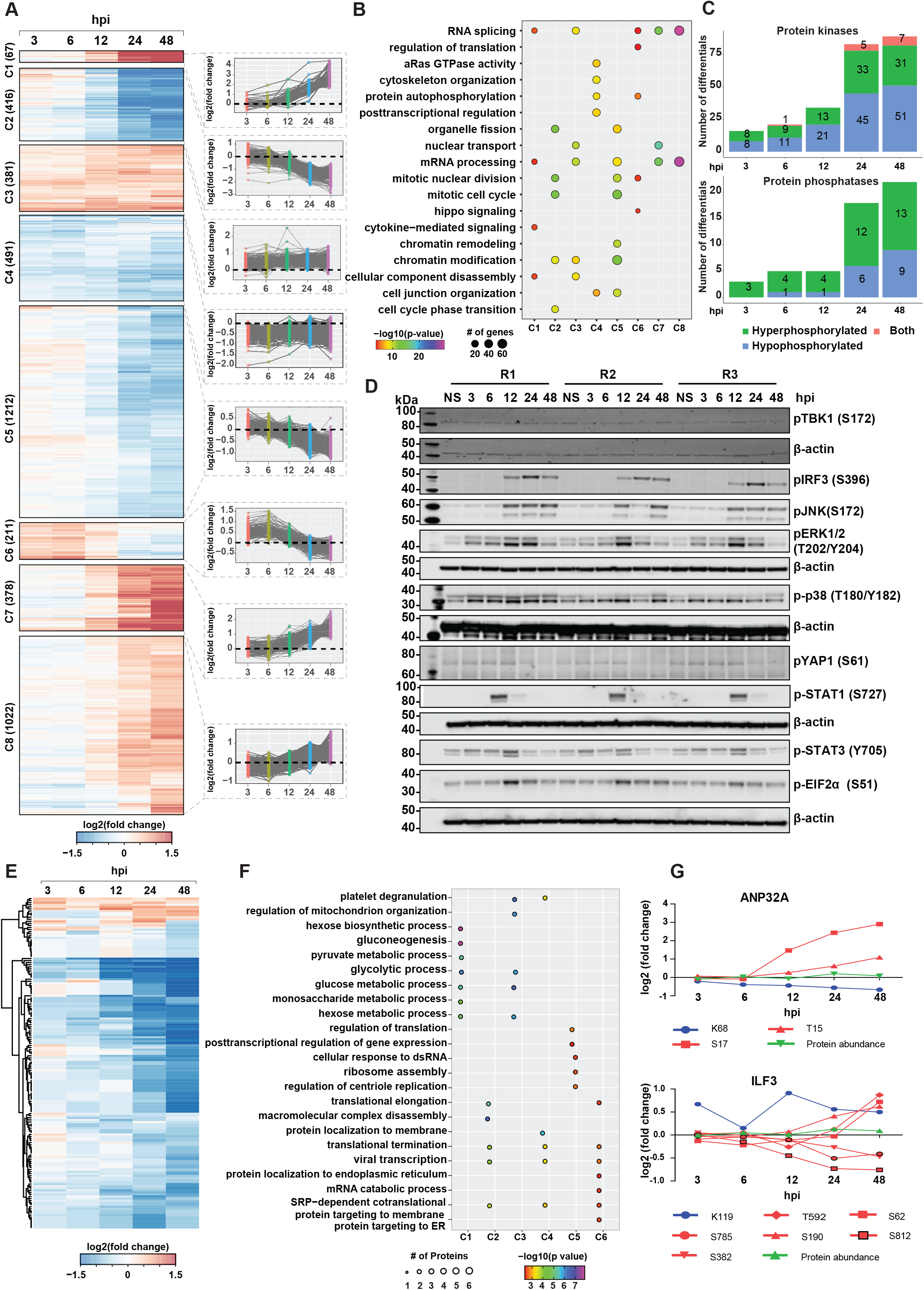
Phosphoproteomics and acetylomics analysis of SARS-CoV-2 infected Calu-3 cells. **A.** Heatmap depicting k-means clustering of phosphoproteomics profile of Calu-3 cells after infection with SARS-CoV-2/ Trondheim-S15/2020 strain at 3, 6, 12, 24, and 48 hpi. **B.** A list of enriched signaling pathways from the phosphoproteomics profile of Calu-3 cells after SARS-CoV-2 infection **C.** Statistics of differentially phosphorylated protein kinases and phosphatases in response to SARS-CoV-2 infection **D.** Immunoblot analysis showing the phosphorylation dynamics of various signaling proteins, including pTBK1 (S172), pIRF3(S396), pJNK(S172), pERK1/2(T202, Y204), p-p38(T180.Y182), pYAP1(S61), p-STAT1(S727), pEIF2α(S51), and p-STAT3 in response to SARS-CoV-2 infection at 3, 6, 12, 24, and 48 hpi. **E.** A heatmap depicting clustering of acetylome profile of Calu-3 cells after infection with SARS-CoV-2/ Trondheim-S15/2020 strain at 3, 6, 12, 24 and 48 hpi. **F.** A list of enriched signaling pathways from the acetylome profile of SARS-CoV-2 infected Calu-3 cells. **G.** Examples of two proteins-ANP32A and ILF3 that are both acetylated and phosphorylated upon SARS-CoV-2 infection.

Notably, differential phosphorylation of signaling proteins related to the antiviral immune response (Cluster 1), including increased phosphorylation of STAT1 and STAT3 at S727 12 hpi, was observed with sustained levels seen even at 48 hpi (**Figure 3A**). Immunoblot analysis further validated our findings (**Figure 3D**). Phosphorylation and activation of STAT1/3 are required for the complete transcriptional activity of the ISGF complex in inducing ISGs and antiviral activity. However, contrary to our findings, Mu *et al*. report that SARS-CoV-2 N protein antagonizes interferon signaling by interacting with STAT1/STAT2 and inhibiting its phosphorylation and nuclear translocation ^60^. Increased phosphorylation of IRF9 at S136 and IRF3 at S396 (**Supplementary Table S3, Figure 3D)** was also observed. IRF9 interacts with phosphorylated STAT1 and IRF3 to form the ISGF3 complex, indicating induction of antiviral immune response that correlates with increased expression of ISGs. These findings strongly establish links to increased ISG levels observed both at the transcript and proteome level, implicating the establishment of a host antiviral state.

Cluster 2 revealed significant enrichment of time-dependent decrease in phosphorylation of proteins involved mainly in the regulation of cell cycle processes, DNA damage and repair pathways and chromatin modification such as CDK1, CDK2 (T14, Y15), CDKN1A (S130), DNA topoisomerases (TOP2A (S1106, S1213, S1504), TOP2B (T1431) and transcription factors/co-regulators- RB1, APC, TP53). Enrichment of DNA damage response pathway at the early time point corroborates with an earlier study that reported that infectious bronchitis virus (IBV) activated ATR-dependent cellular DNA damage response to induce cell cycle arrest for the enhancement of viral replication and progeny production. They further demonstrated that the interaction between *Coronavirus* nsp13 and DNA polymerase δ was essential for the induction of DNA damage response and cell cycle arrest at the S phase ^61^. Responses enriched as early as 3hpi indicate enrichment of protein autophosphorylation required for activation of kinases modulating downstream cellular processes.

Hypophosphorylation of several sites on proteins that regulate actin cytoskeleton organization (cluster 4) was observed upon SARS-CoV-2 infection. It has been previously shown that sustained RAF/MAPK signaling results in the downregulation of ROCKI and Rho-kinase, two-Rho effectors required for stress fiber formation and promote cytoskeleton reorganization ^62^. Inhibition of MEK functionally restored the activity of ROCK1/Rho-kinase in promoting cytoskeleton reorganization in NRK/RAS cells. We observed sustained MAPK1 activation upon SARS-CoV-2 infection with decreased levels of pROCK1 24 hpi, thereby suggesting potential alterations in cytoskeleton organization. Increased phosphoERK1/2 (T202, Y204) was confirmed by immunoblotting (**Figure 3D**). Clusters C2 and C6 show a significant time-dependent decrease in phosphorylation, specifically at 24 hpi. Proteins involved in Hippo/YAP signaling, Signaling by Rho GTPases, vesicle-mediated transport, tight junction proteins, including TJP1, TJP2, CTTN, SYMPK, among others, were significantly enriched. Hippo/YAP1 Signaling was enriched specifically in cluster 6, corroborating with hypophosphorylation of YAP1, an essential component of the Hippo/YAP1 signaling pathway, at S61. We also observed hypophosphorylation of other Hippo signaling proteins - TJP1, TJP2, DLG1, SCIB, among others. A time-dependent decrease in pYAP1 (S61) was independently confirmed by immunoblotting **(Figure 3D)**. Hippo signaling is further discussed in subsequent sections.

We also observed temporal changes in the SARS-CoV-2-responsive kinome and phosphatome (**Figure 3C**), with several protein kinases demonstrating decreased phosphorylation levels over time. On the contrary, the phosphorylation levels of protein phosphatases such as PTPN3, CTDSPL, and CTDSPL2 increased with time, with 9 phosphatases hyperphosphorylated at 24 hpi. Of the 107 kinases found to be differentially regulated, 18 have been described as substrates of viral proteins ^23, 29^, including kinases involved in PI3K/AKT signaling (GSK3B, RAF1, PAK4), cell cycle (PRKDC, CDK2, and TTK) and MARK kinase signaling (MARK2 and MARK3). Further PTM signature enrichment analysis using PTMSigDB revealed enrichment of signatures of MAPK signaling pathways including MAPK14, MAPKAP2, leptin pathway, PRKACA, cell cycle kinases CDK1, CDK2, and Aurora B at early time points of infection correlating with activation of DNA damage and repair pathways and altered cell cycle regulation (**Figure S3B**). Furthermore, a substantial number of transcription factors were differentially phosphorylated in response to SARS-CoV-2 infection (**Figure S3C and S3D**). Notably, TAF6 (S672) and TAF7 (S264), components of the DNA-binding general transcription factor complex TFIID, were hypophosphorylated upon SARS-CoV-2 infection, especially TAF7, as early as 6 hpi. Phosphorylation of TAF7 (S264) mediated by TAF1 has been shown to influence the levels of cyclin D1 and cyclin A gene transcription by increasing TAF1 histone acetyltransferase (HAT) activity and histone H3 acetylation levels ^63^.

Emerging evidence strongly indicates the role of protein lysine acetylation as a regulatory mechanism in viral infection and, therefore, can likely serve as potential therapeutic targets. However, the dynamic alterations in protein acetylation upon SARS-CoV-2 infection have not been explored. Analysis of protein lysine acetylation dynamics revealed 112 acetylated peptides differentially regulated upon SARS-CoV-2 infection (**Figure 3E**). Of these, sitespecific alterations in acetylation of 53 proteins were largely independent of protein abundance changes except for CDK1(K33), which was found to be hypoacetylated at 24 hpi. In contrast, the protein expression was downregulated only at 48 hpi. On the contrary, the acetylation levels of TMSB4X at K26 and K39 decreased at 6 hpi with the proteome expression downregulated 24 hpi *k*-means clustering of the acetylome measurements enabled the segregation of the regulated genes into early, late, and mid-late responders to viral infection. We largely observed significant downregulation of acetyl sites on proteins as early as 6 hpi with respect to Clusters 2 and 3. In contrast, Cluster 1 included acetylation sites on proteins that were predominantly downregulated at a late time-point (48 hpi). Clusters 4 and 5 revealed hypoacetylation at 6 hpi in comparison with 3 hpi followed by increased acetylation (log2FC>0.5) post 12 hpi

Pathway enrichment analysis largely revealed the enrichment of proteins involved in various biological processes (**Figure 3F)**. Notably, a time-dependent decrease in the acetylation levels across proteins belonging to several classes but primarily histones, epigenetic modifiers, and proteins involved in metabolic regulation. Cluster 1, which largely showed a delayed response, was enriched in proteins involved in glucose metabolism and hexose biosynthetic process, as well as histone subunits (H2B and H4C), acetylated on multiple sites. In addition to proteins enriched in glucose/hexose metabolism, Cluster 3 consisted of hypoacetylated proteins involved in the regulation of the mitochondrial organization. Acetylation of GAPDH at K251 was observed as early as 3 hpi, followed by a persistent decline at later time points of infection. Acetylation at this site is known to be mediated by PCAF and is required for nuclear translocation of GAPDH during apoptotic stress ^64^. In the case of other metabolic enzymes, a progressive decline in the acetylation status was observed across time points, with a significant reduction post 24 hours. Cluster 2 and Cluster 4 were enriched in proteins involved in viral transcription and translation termination, further corroborating the role of viruses inducing host protein translation shutoff activity. Additionally, proteins localized to the membrane were enriched in Cluster 4, whereas proteins involved in macromolecule complex disassembly and translational elongation were enriched in Cluster 2. Acetylation of EIF5A2, a translation initiation factor essential for cell growth at K47, is responsible for regulating its subcellular localization with the hypoacetylated form predominantly localized to the cytoplasm ^65^. Cluster 5 includes histones-H2AZ1 acetylated at K8, K12, and K14, H3C1 (K24), as well as keratins KRT17 (K219) and KRT19 (K215) were found to be hyperacetylated 12 hpi. H3C1 is a core component of the nucleosome and is involved in the process of post-transcriptional and translational regulation of genes. Furthermore, studies have revealed that this site is amenable to acyl modifications and is highly responsive and reversibly regulated by nutrient availability ^66^. Overall, our data is indicative of SARS-CoV-2 mediated triggering of deacetylases that results in significant protein deacetylation at later time points of infection. These results correlate with an earlier report suggesting that active deacetylase activity is required to induce ISG expression and antiviral immune responses ^67^, thereby creating an environment conducive to increased viral replication.

We next assessed if there was an overlap of dysregulated phosphorylation and acetylation datasets. Our data clearly shows an interplay of phosphorylation and acetylation on host proteins. In all, we identified 37 proteins with differentially regulated multiple PTM sites. These include proteins involved in innate immune signaling (PPIA, HSP90AA1, ILF3, ANXA2, ANXA4, and CFL1), regulation of cellular metabolic processes as well as proteins involved in the regulation of mRNA stability, including ANP32A, a multifunctional transcriptional regulator and a component of the inhibitor of histone acetyltransferases complex. Significant hypoacetylation was observed at 24-48 hpi, with the phosphorylation levels on T15 and S17 showing an opposite trend (**Figure 3G**). Interestingly, the levels of ubiquitination at K99, as demonstrated by Stukalov *et al*., were downregulated as early as 6 hpi with a minimal increase over time. Host proteins, especially ANP32A, interact with influenza virus RNA polymerase components, forming a replication platform essential to promoting vRNA synthesis^68, 69^. Further, ANP32A is known to be a component of the inhibitor of histone acetyltransferases complex and modulates the HAT activity of EP300/CREBBP (CREB-binding protein) and EP300/CREBBP-associated factor ^70, 71^ and is involved in the positive regulation of ISG transcription ^72^ which we observe at both transcript and protein levels. It remains to be determined if the differential acetylation at this site located in the LRR domain plays a role in CoV-2 replication and infectivity. As described previously, we also identified distinct phosphorylation and acetylation patterns on several sites on Vimentin (VIM), an essential host factor responsible for the entry and pathogenesis of SARS-CoV-2 ^73^. Decreased acetylation levels as early as 6 hpi (K294, K402) whereas distinct phosphorylation on several sites early (S73) or late (S83, S325, S420, and S430) were observed (**Figure S3E**). Interestingly, Stukalov *et al*. report decreased ubiquitination levels following infection with SARS-CoV-2 and SARS-CoV.

A comparison of the changes in the acetylation data with the ubiquitinome performed in ACE2-A549 cells ^29^ revealed 26 proteins common across the data sets. Of these, six proteins, including ribosomal proteins-RPL13A (K159), RPL34 (K36), RPL7 (K29), RPS25 (K57); LRRC59 (K135), and UBE2N (K82) were differentially modified at the same residue. However, the PTM dynamics observed were opposite in trend, especially at 24 hpi, wherein the acetylation levels were downregulated as opposed to increased ubiquitination. Notably, LRRC59, a leucine-rich repeat-containing ER membrane protein, plays an important role in innate immune signaling by modulating DDX58-mediated type I IFN signaling ^74^ and regulating trafficking of nucleic-acid sensing TLRs ^75^ was found to be hypoacetylated at 6 hpi with acetylation reaching a basal level at 12 hpi. On the contrary, the ubiquitination levels at K135 and others (K71, K207) remain unchanged at 6 hpi but gradually increase across time points, with significantly elevated levels observed at 24 hpi (**Figure S3F**). Overall, our findings further emphasize the need to assess the effects of different post-translational modifications simultaneously, as the functional outcome of the cellular response is dependent on the concerted action of regulatory machinery.

### 4. Modulation of secretory effectors involved in SARS-CoV-2 infection

To identify molecular/metabolic pathways perturbed following SARS-CoV-2 exposure, we measured the extent of release of 100 metabolites in the culture supernatants of Calu-3 cells treated with SARS-CoV-2 for the time points described above (**Figure 1A**). Of these, 85 were identified and quantitated across five time points after SARS-CoV-2 infection and were considered for further analysis. A total of 163 differential events were identified across the five time points, of which 146 were increased, and 17 were decreased (**Figure 4A**). The metabolites that were significantly changed in response to SARS-CoV-2 infection have been provided in **Supplementary Table S6**.

**Figure 4.**
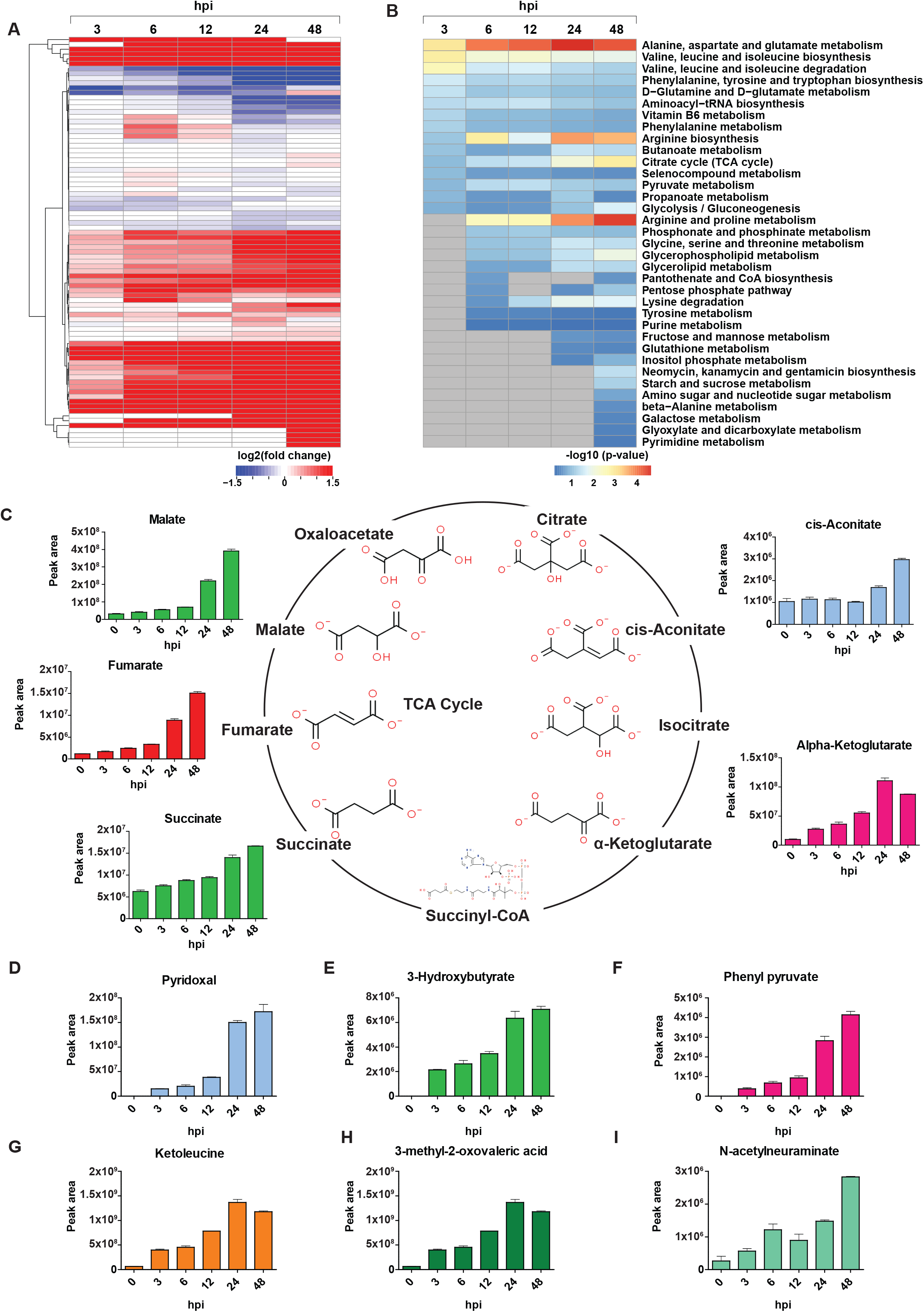
Metabolomics analysis of SARS-CoV-2 infected Calu-3 cells. **A.** A heatmap depicting differential levels of metabolites in response to infection with SARS-CoV-2/ Trondheim-S15/2020 strain at 3, 6, 12, 24, and 48 hpi. **B.** A list of enriched metabolic pathways from the metabolomics profile of Calu-3 cells upon SARS-CoV-2 infection **C.** A schematic depicting the TCA cycle and differential levels of TCA metabolites-cis-aconitate, alpha-ketoglutarate, succinate, fumarate, and malate in response to SARS-CoV-2 infection. Altered levels of metabolites, including **D.** Pyridoxal, **E.** 3-hydroxybutyrate, **F.** Phenylpyruvate, **G.** Ketoleucine, **H.** 3-methyl-2-oxovaleric acid, and **I.** N-acetylneuraminate observed in response to SARS-CoV-2 infection.

Changes in the metabolite release and metabolic pathway perturbations were analyzed using targeted metabolome-wide association. Metabolite set enrichment analysis (MSEA) indicated significant enrichment of several metabolic pathways, including the TCA cycle, Glycolysis/Gluconeogenesis, butanoate metabolism, and metabolism pathways of several amino acids (**Figure 4B, Figure S4A-E)**. Metabolites involved in Amino acid metabolism, including branched-chain amino acids, and aromatic amino acids, were among the significantly enriched pathways at early and mid-time points of infection. At later time points, arginine metabolism and metabolites of the TCA cycle pathways were found to be significantly enriched. However, the extent of changes in the levels of arginine was not significant across the time points. Intermediates of the TCA cycle were found to show altered levels in response to SARS-CoV-2 infection (**Figure 4C**). An increase in succinate levels was observed at 24-48 hpi, while fumaric acid levels increased at 6-48 hpi, and alpha-ketoglutarate levels were increased across all the studied time points. However, cis-aconitate was only increased at 48 hpi while pyruvic acid levels decreased at 12-48 hpi. Several metabolites showed increased levels across the studied time points (**Figure 4D-I**). These include the vitamin Pyridoxal, Phenylpyruvate, organic acids such as lactate, Methylmalonic acid, and branched-chain amino acid metabolism intermediates: 3-methyl-2-oxovaleric acid and Ketoleucine and 3-Hydroxybutyrate, also known as Beta-hydroxybutyrate (BHB). This endogenous ketone body acts as a highly efficient oxidative fuel. Recent studies implicate its vital role in immunomodulation ^76^, Interestingly, we observed increased levels of sialic acid N-Acetylneuraminate as early as 3 hpi, and it has been recently demonstrated that N-acetylneuraminic acid (Neu5Ac) serves as a plausible alternative receptor for SARS-CoV as a key domain in CoV-2 spike protein binds to Neu5Ac, a process essential for viral entry into cells ^77, 78^. The increased secretion of several acids and ketone bodies in response to SARS-CoV-2 infection at all time points suggests metabolic ketoacidosis.

### 5. Global alterations in the viral proteome profile

In addition to the changes in host cellular proteins, analysis of upregulated DEGs revealed a significant increase in the expression of viral genes observed mainly at 24 hpi (**Supplementary Table S7**). This was demonstrated by the low alignment rate observed against the human reference database, indicating that the RNA fraction at these time points was taken over by the virus. At the proteome level, nine canonical SARS-CoV-2 proteins were consistently identified and quantified across six time points of infection. In concordance with previous reports, over 3-fold induction of viral proteins with similar expression trends across all nine viral proteins was observed (**Figure 1C**). The dynamics of protein expression across time points indicated that the viral protein synthesis increased continuously post-infection, with a peak observed at 24 hpi except for Orf8, with a slight decrease in expression observed at 48 hpi (**Figure 1C, Supplementary Table S2**). SARS-CoV-2 N, M, S, and Orf9b were among the most abundant proteins consistently detected.

Analysis of the PTM dynamics revealed 41 phosphorylation sites on eight viral proteins with maximum changes in the phosphorylation dynamics observed post 12 hpi (**Figure 5A**). The trend observed was in accordance with the increase in protein abundance. Comparison with previously reported SARS-CoV-2 phosphoproteome enabled us to identify ten novel phosphorylation sites on 6 viral proteins, including ORF1a (S142, S2517) and ORF3a (T24, S272), ORF7a (S44), ORF8 (S103) (**Figure 5B**). Replicase protein 1a (P0DTC1) is a polyprotein that is proteolytically cleaved to generate several non-structural proteins. We found the polyprotein to be modified on 3 sites; one corresponding to NSP1 (S142) and 2 sites corresponding to NSP3 (S2517, S2644). In concordance with earlier studies, multiple sites of phosphorylation on the nucleocapsid protein, including 3 novel sites corresponding to T245, S327, and S379, clustered in the linker region between the RNA-binding (RBD) and dimerization domains as well as located in the dimerization domain (S327) were observed (**Figure 5C**). The nucleocapsid protein plays an essential role in the replication, transcription, and assembly of the SARS-CoV-2 genome ^79^ and is also known to control host cell cycle machinery ^23, 80^. The identified phosphorylation motifs were bioinformatically assessed using NetPhos 3.1 to predict potential host kinases phosphorylating the viral proteins (**Figure 5D**). Consistent with previous findings, we too observed several motifs in nucleoprotein to be modified by CMGC kinases, including members of MAPK family CDKs and GSK3. Interestingly, we observed several novel phosphorylation sites (N, ORF1a, ORF8) mapping to recognition motifs of the AGC family -PKA, PKB, and PKC and DNAPK, a critical player involved in the DNA damage response pathways. It is known that DNA-PK acts as a DNA sensor that activates innate immunity. However, its role in phosphorylating RNA virus N protein, especially at the novel site T379 located C-terminal of the dimerization domain, remains to be explored ^81^. In concordance with previous results, we also observed an increased phosphorylation level of nucleocapsid protein. A recent study observed the highest percentage of disorder compared to other viral proteins as well as a high number of variable molecular recognition features (MoRFs) and classified as a highly disordered protein with a central role in viral pathogenesis. Furthermore, the viral acetylome profile revealed hyperacetylation of 6 sites on nucleoprotein, including at K65, K248, K249, K256, K341, and K362. Of these, two sites on the nucleoprotein (K248, and K249) have been recently demonstrated to be acetylated by P300/CBP-associated factor (PCAF) and general control nonderepressible 5 (GCN5) ^82^. The hyperacetylation of N protein coincides with the hypoacetylation of ANP32A and the induction of ISGs, which strongly demonstrates the critical role of ANP32A in modulating the host PTM dynamics, thereby contributing to viral pathogenesis.

**Figure 5.**
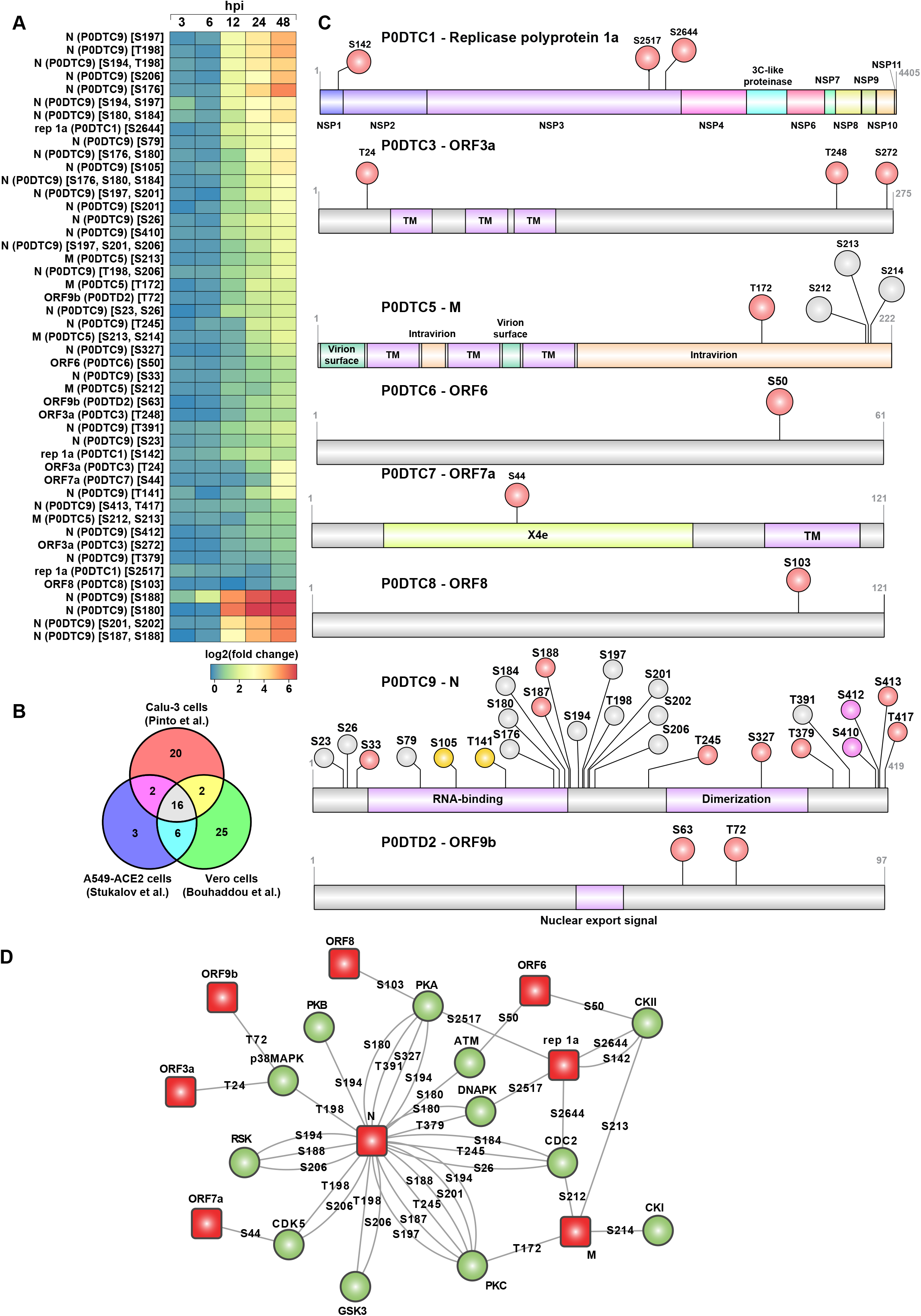
Viral phosphoproteomic dynamics after infection of Calu-3 cells with SARS-CoV-2/ Trondheim-S15/2020 strain. **A.** Heatmap revealing temporal phosphoproteomic changes in SARS-CoV-2 proteins **B.** Comparison of viral phosphoproteome identified in the current study with phosphoproteomics data from previously published studies. **C.** Schematic shows phosphosites’ overlap on important domains and motifs in SARS-CoV-2 viral proteins. **D.** Upstream kinase analysis using NetPhos 3.1 to predict potential host kinases phosphorylating the viral proteins. Red boxes represent viral proteins, and the green circles represent host kinases. The phosphorylated residue on each viral protein is indicated.

We next looked at the positions of phosphorylated residues in the available and predicted structural models of SARS-CoV-2 proteins (**Figure S5**). For those residues that could be mapped on the available structures, all the identified phosphorylation sites were plausible since they were located at the surface of the proteins and, therefore, should be accessible to kinases. Similar to the previous reports ^20^, most of these residues were located in loops or at the edge of secondary structure elements. The majority of the side chains were not engaged in any intramolecular interactions (for example, S63 of ORF9b or T141 and S327 of N protein), though some residues formed hydrogen bonds that could support the positioning of some flexible loops (for example, S103 of ORF8 or S44 of ORF7a) or contribute to an interaction with another protein (T72 of ORF9b bound to the backbone of V556 of Tom70). In addition, many phosphorylation sites were found in regions that were missing from crystal or cryoelectron microscopy structures and therefore presumed to be likely in unstructured regions (for example, T24, S272, and T248 of ORF3a). Indeed, one such region was S176-S180 of Protein N, which was modeled as a flexible linker in the NMR ensemble ^83^ and is part of the serine/arginine-rich (SR) domain at the start of one of the predicted intrinsically disordered regions (IDR) (residues 1-44, 175-254, 365-~400). The SR region may participate in RNA binding ^83^ and is a hotspot for mutations in Protein N ^84^. Interestingly, S327 of N Protein, a phosphorylation site unique to our study, has also been found to be frequently mutated, typically to Leu ^84^. For some of the most interesting cases for which experimental models were missing, we used computational modeling with TrRosetta ^85^. Although computational models should be approached as only approximate, especially in the positioning of flexible regions and the orientation of independent domains, they all showed that the phosphorylation sites were located on protein surfaces and in flexible regions, including T172, S212, and

S213 of Protein M, T24, S272 and T248 of ORF3a, S50 of ORF6, S26, T245 and T379 of Protein N, S142 of Rep1a (Nsp1), S2517 and S2644 of Rep1a (Nsp3). In summary, the phosphorylation sites observed in this study were predicted to be unlikely to play an important structural role in supporting the architecture of SARS-CoV-2 proteins, which corresponded well to their exposed positions and the lack of conservation noted in some cases. Instead, phosphorylation might play a role in regulating interactions mediated by these residues – the change in electrostatic potential or in the pattern of hydrogen bonds could alter the binding of other proteins or RNA. Whether these could be beneficial or detrimental to the SARS-CoV-2 remains to be validated by functional studies.

### 6. Integrated analysis of multi-omics datasets identified aberrant Hippo signaling, DNA damage/repair, and ubiquitination machinery of host cells

Integrated analysis of key signaling pathways and processes identified in our datasets using pre-defined gene sets from MSigDB revealed significant changes at multiple levels on proteins involved in Hippo signaling, DNA/damage/response, protein ubiquitination, and alternate splicing pathways (**Figure 6A-G, Figure S6A-H**). While the phosphoproteome data showed a time-dependent decrease in the phosphorylation status of Hippo signaling, other pathways, including DNA damage/response, protein ubiquitination, and spliceosome pathways, demonstrated increased signaling. The proteome changes showed reduced expression, while the transcriptome showed increased transcription of genes belonging to these pathways.

**Figure 6.**
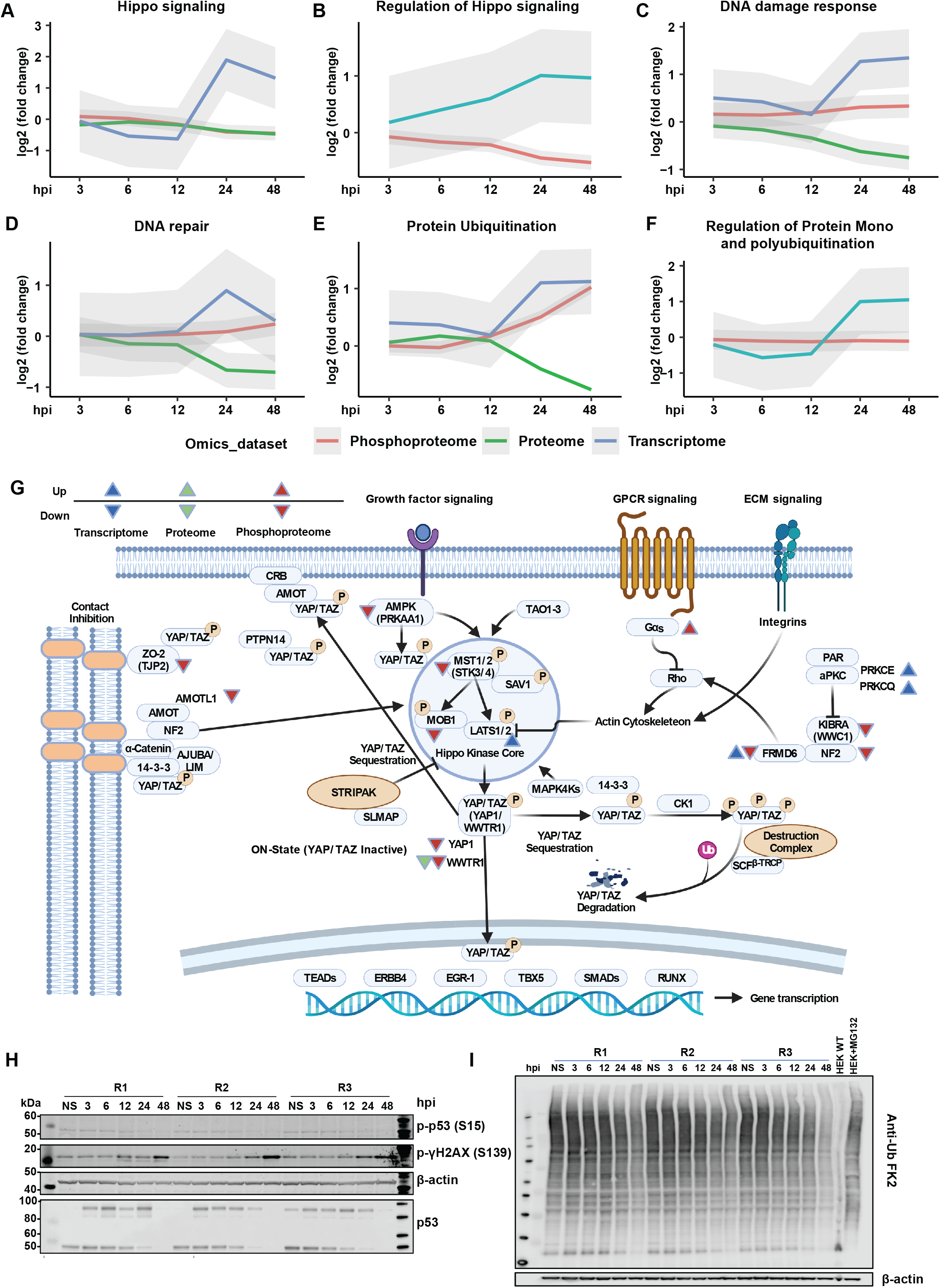
Integrated omics analysis identifies alterations in key signaling pathways upon SARS-CoV-2 infection in Calu-3 cells. Graphs illustrating the average trend of differentials from transcriptomics, proteomics, and phosphoproteomics datasets with respect to various pathways and processes, including **A.** Hippo signaling **B.** Regulation of Hippo signaling, **C.** DNA damage response, **D.** DNA repair, **E.** Protein ubiquitination, **F.** Regulation of protein mono and polyubiquitination. **G.** A detailed overview of the Hippo signaling pathway indicating differentials in response to SARS-CoV-2 infection. **H.** Immunoblot analysis of DNA damage markers, including phospho-p53 (S15), and phospho-γH2AX (S139) along with total p53 and β-actin (control) in Calu-3 cells in response to SARS-CoV-2 infection at 3, 6, 12, 24, and 48 hpi. **I.** Immunoblot analysis indicating changes in the total protein ubiquitination profile of Calu-3 cells in response to SARS-CoV-2 infection at 3, 6, 12, 24, and 48 hpi.

While in cancers, the Hippo signaling pathway is a tumor suppressor by nature ^86^, various studies have indicated its crucial role in viral infections such as those caused by Hepatitis B virus (HBV), Hepatitis C virus (HCV), Human papillomavirus (HPV), Epstein–Barr virus (EBV), Zika virus (ZIKV), amongst others ^87^. During the course of Hippo signaling, MST1/2 kinases (STK3/STK4) phosphorylate and activate Lats1/2 kinases, which in turn phosphorylate and inhibit essential transcriptional coactivators -YAP/TAZ (YAP1/WWTR1) ^86, 88^. When upstream kinases are repressed, dephosphorylated YAP/TAZ accumulate in the cell nucleus and associate with several transcriptional factors that stimulate genes involved in cell survival and proliferation, including TEADs, ERBB4, EGR1, TBX5, and members of the SMAD and RUNX family ^89^. Notably, YAP was found to negatively regulate the antiviral immune responses in Sendai virus (SeV), vesicular stomatitis virus (VSV), and herpes simplex virus type 1 (HSV-1) infections ^90^. Wang and colleagues further demonstrated that depletion of YAP in macrophages increased interferon beta expression. Also, YAP overexpression led to repression of IRF3 dimerization through YAP association and decreased nuclear localization of IRF3. Further, viral-mediated activation of IKKε phosphorylated YAP led to its degradation, thereby relieving cells of YAP-mediated antiviral response. Our phosphoproteomics experiments observed the altered phosphorylation of Hippo signaling-related proteins in response to SARS-CoV-2 infection. Recently a preprint by Garcia et al. showed that activation of the Hippo signaling pathway during SARS-CoV-2 infection contributed to host antiviral response ^91^. Further, they used a pharmacological inhibitor of YAP, significantly reducing SARS-CoV-2 infection. This study corroborates our findings on the importance of Hippo signaling in the context of SARS-CoV-2 infection. In the current study, we observed hypophosphorylation of upstream kinases involved in Hippo signaling, including STK3 (S386, T354, S370/S371), and STK4 (S40, T41, S43). Further, we observed YAP1 (S61) and WWTR1 (S62, T67) phosphorylation levels progressively decreasing and the lowest at 48 hpi. Additionally, validation using immunoblot corroborated the hypophosphorylation of YAP1 at S61. Further, we also observed hypophosphorylation of PRKAA1 or AMPK (S486, T490), which is known to phosphorylate YAP1 at S61 ^92^. We also found other regulators of the Hippo signaling pathway such as KIBRA or WWC1 (S947), FRMD6 (S375), NF2 (S518), and ZO-2 or TJP2 (S966, S1027) to be hypophosphorylated. Signaling crosstalk between the Hippo and TGFβ pathway is known ^93^. In our study, TGFβ receptor TGFBR2 was progressively hyperphosphorylated, reaching maximal hyperphosphorylation at 48 hpi. LATS-mediated phosphorylation of YAP results in its interaction and binding to 14-3-3, leading to cytoplasmic retention ^94^. We identified hypoacetylation of both YWHAB (14-3-3β) at K13 and YWHAZ (14-3-3ζ) at K11. Whether altered acetylation directly affects YAP localization and activity remains to be determined. While it is well known that metabolic cues can control YAP/TAZ regulation ^95, 96^, increasing evidence suggests YAP influences cellular metabolism. YAP has been demonstrated as a central regulator of glucose metabolism ^97^, mediating the regulation of glucose transporter 1 (glut1) in zebrafish. Yap mutant zebrafish were reported to have impaired glucose uptake, nucleotide synthesis and glucose tolerance in adults ^98^. Our study indicates high levels of glycolysis intermediates, including glucose 6-phosphate, fructose 6-phosphate, dihydroxyacetone phosphate, and lactate at 48 hpi. These findings likely suggest that YAP and, thereby, hippo signaling could profoundly impact glucose metabolism during SARS-CoV-2 infection. We further observed increased alpha-ketoglutarate, proline, and uracil levels in response to SARS-CoV-2 infection and likely associate this observation with dysregulated YAP/Hippo signaling as YAP has been found previously to regulate fatty acid oxidation and amino acid metabolism ^99^. YAP inhibition in the skeletal muscles decreased levels of various metabolites, including undecanoic acid, capric acid, 2-octanoic acid, 2-oxoglutaric acid (alpha-ketoglutarate), amino acids-lysine, serine, proline, aspartate and uracil intermediate, 3-ureidopropionic acid. YAP has also been shown to regulate glutamine metabolism to stimulate nucleotide biosynthesis through increased expression of glutamine synthetase in zebrafish ^100^. In light of the dysregulated metabolites and aberrant Hippo signaling identified in the current study in response to SARS-CoV-2 infection, further investigation into the possible influence of Hippo signaling on cellular metabolism during viral infection is warranted.

Further, the DNA damage response pathway was enriched across multiple omics datasets in response to SARS-CoV-2 infection. We identified p53 S314/315, not S15, and the phosphorylation level was downregulated (log_2_FC −0.5). Immunoblot analysis revealed decreased phosphorylation of p53 S15 and corresponded to the reduced abundance of total p53 levels indicating CoV-2 infection potentially mediates p53 degradation (**Figure 6H**). At later time points of infection, we observed hyperphosphorylation of Ser/Thr kinase PRKDC (also known as DNAPK) at multiple sites. DNAPK acts as a molecular sensor for DNA damage and is also known to regulate DNA virus-mediated innate immune response. We identified hyperphosphorylation of several DNAPK sites including S3205, an ATM target phosphorylated either through autophosphorylation ^101^ or through PLK1 ^102^ and S2672, which is part of the ABCDE cluster, which promotes DNA end processing, and autophosphorylation of this cluster is required to initiate DNA damage repair ^101^. Regulation of DNA damage response mechanism by DNAPK is mediated by phosphorylation of S139/S140 of histone variant H2AX/H2AFX. Concurrently, we observed increased phosphorylation of γH2AX at S140 from 12 hpi with no changes in the total protein levels. We further confirmed our results through immunoblot analysis (**Figure 6H**).YAP1/Hippo signaling has also been known to cross-talk with the DNA damage response pathways ^103, 104, 105, 106, 107^, and this could help explain why DNA damage/repair pathways were enriched in Calu-3 omics data after SARS-CoV-2 infection. Further, YAP and P53 have been known to crosstalk via a SIRT-1 mediated deacetylation mechanism, thereby controlling the cell cycle in A549 lung cells ^108^. TAZ (or WWTR1) has also been known to reduce P53 activity through p300-mediated acetylation negatively 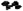. Finally, ANP32E has previously been reported to remove H2A.Z from DNA double-strand breaks, promoting DNA repair and nucleosome organization ^109^. All these findings suggest a deeper connection between DNA damage and YAP/TAZ signaling.

Ubiquitination is essential in regulating the innate immune response. E3 ligases, such as TRIM25 and RIPLET, have been shown to mediate RIG-I ubiquitination and type I IFN induction^107^. We observed several Ubiquitin (Ub) E2 and E3 ligases such as UBE2C, UBE2J1, UBE2S, PJA2, RNF5, RNF14, and RNF115, among others, to be significantly downregulated as early as 12 hpi (**Figure S6I**). To assess if this affected the status of protein ubiquitination, the pan-Ub profile was explored using Anti-Ub FK2, which detects K29-, K48-, and K63-linked mono- and polyubiquitinylated proteins. Similar to the expression profiles of E3 ligases, a significant decrease in ubiquitination was observed, especially at 24 and 48 hpi (**Figure 6I**), suggesting CoV-2-mediated hijacking of the host ubiquitin system to maximize their own survival likely through mechanisms adopted by DNA tumor viruses and Adenoviruses. A recent study aimed to identify substrates of the Mpro from three coronaviruses revealed many E3 ubiquitin ligases were cleaved by the SARS-CoV-2 main protease Mpros including RNF20, ITCH, UBE3A, confirming our findings that albeit increased expression at the transcript level, profound downregulation of E3 ligases owing to degradation was essential to counteract the host innate immune response ^107, 110^. Additionally, Coronaviruses encode papain-like protease (PLP) that acts as a cysteine protease as well as possesses intrinsic deubiquitinating and deISGylating activities required for viral replication and the evasion of host responses.

We observed time-dependent dephosphorylation of several proteins involved in the cell cycle at multiple sites, including Rb, which acts as a key driver of cell cycle regulation. A time-dependent decrease in phosphorylation at S249, T821, and T826 is known to be mediated by members of type 1 serine/threonine protein phosphatases (PP1) ^111^ was observed. Earlier studies indicate that endogenous Rb is dephosphorylated on Threonine-821 when cells undergo apoptosis ^112^. Changes in the phosphorylation dynamics were accompanied by downregulation at the proteome level for most proteins involved in regulating cell cycle processes. (**Figure S6J**).

## Conclusions

We describe a systems-level approach to discover signaling pathways modulated upon infection of Calu-3 cells with SARS-CoV-2 S15 Trondheim strain belonging to the wave 1 isolates. In-depth systems-wide and time-resolved characterization of the host and viral changes throughout productive infection revealed temporal changes in the transcriptome, proteome, phosphoproteome, acetylome, and exometabolome. While other studies have extensively explored SARS-CoV-2 mediated alterations in the proteome and phosphoproteome of infected cells ^27, 29, 31, 44, 45^, to our knowledge, this study is the first to perform comprehensive profiling, including exploring the acetylome and exometablome dynamics in response to SARS-CoV-2 infection. Multiple genes and functional pathways identified in our data were previously reported to promote SARS-CoV-2 mediated pathogenesis, validating the rigor of our approach and providing further support for the role of these specific host factors. In concordance with earlier studies, we too observed a robust induction of the innate immune response accompanied by increased interferon signaling, increased expression of ISGs, TLRs, and cytokines and chemokines. Further, our results are concordant with studies showing that the pulmonary viral load positively correlates with the expression of ISGs, with reduced risk and disease severity ^113, 114, 115^. Integrated analysis revealed changes in Hippo signaling, DNA damage response, ubiquitination, and spliceosome machinery pathways across omics datasets. In response to infection, several novel phosphorylations and acetylation sites were identified on vital host proteins. We also provide evidence of host-mediated PTM modification of viral proteins, including time-dependent alterations in the acetylation levels of nucleoprotein for the first time. Altered PTM levels have been demonstrated to influence virus-mediated control of host cellular dynamics and the role of novel acetylation sites warrants further investigations. Altered release of TCA cycle intermediates with increased secretion of organic acids and ketone bodies was observed and could likely be a counteractive measure to restrict SARS-CoV2 infection^116^. Finally, the data from the study shows evidence of potential crosstalk between Hippo signaling, DNA damage response, and post-translational modifications. However, the site-specific alterations in the PTM dynamics and it effect on the signaling crosstalks in response to SARS-CoV2 infection will need to be deciphered in greater detail by future investigations.

Although the data from this study and several published multi-omics studies on SARS-COV-2 infection ^20, 27, 29, 31, 44^ reveal differences that likely arise from the use of different cell types, protocols used for infection, sample preparation and data acquisition strategies, a compilation of the information provides important information on the discreet molecular level regulation. Specifically, the measurement of PTM dynamics across the time course of infection, can be leveraged to investigate immune evasion mechanisms and enhanced transmission. Furthermore, the findings from this study and other studies could serve as a resource for future investigations on newer SARS-COV-2 strains and a point of reference during the evolution of existing strains. Such an approach could aid in future pandemic preparedness in terms of finding novel drug targets.

## Online methods

### Cell lines and reagents

Non-small-cell human lung cancer Calu-3 obtained from ATCC (HTB-55) were grown in DMEM-F12 medium supplemented with 100 μg/mL streptomycin and 100 U/mL penicillin (Pen/Strep), 2 mM L-glutamine, 10% FBS, and 0.25% sodium bicarbonate (Sigma-Aldrich, St. Louis, USA) (complete media). Kidney epithelial cells extracted from an African green monkey (Vero-E6) were grown in DMEM medium supplemented with 10% FBS and Pen-Strep. The cell lines were maintained at 37°C with 5% CO2.

### Virus strains, stock preparation, plaque assay, and in vitro infection

All experiments involving live SARS-CoV-2 were performed in compliance with NTNU’s guidelines for Biosafety Level 3 (BSL-3) containment procedures in approved laboratories. All experiments were performed in at least three biologically independent samples. hCoV-19/Norway/Trondheim-S15/2020 strain from our previous study ^43^ were amplified in a monolayer of Vero-E6 cells in DMEM media containing Pen/Strep and 0.2% bovine serum albumin. The media from the viral culture were serially diluted from 10^-2^ to 10^-7^ in serum-free media containing 0.2% bovine serum albumin (BSA). The dilutions were applied to a monolayer of Vero-E6 cells in 24-well plates. After one hour, cells were overlaid with the virus growth medium containing 1% carboxymethyl cellulose and incubated for 72 hours. The cells were fixed and stained with crystal violet dye, and the plaques were calculated in each well and expressed as plaque-forming units per ml (pfu/mL).

### SARS-CoV-2 infection and cell culture

Calu-3 cells were grown in complete DMEM-F12 media in a humidified incubator at 37 °C in the presence of 5% CO2. The medium was replaced with DMEM-F12 containing 0.2% BSA and Pen-Strep prior to infection experiments. The cells were mock- or virus-infected at an MOI of 0.1 for 0, 3, 6, 12, 24, and 48 hours. At each time point, the samples were washed twice with 1x TBS buffer and harvested in QIAzol lysis reagent (Qiagen, Germany) (for RNASeq) followed by DNAse digestion (Qiagen, Germany) treatment on the RNeasy Mini columns, according to the manufacturer’s protocol. SDS lysis buffer 4%SDS, 50 mM TEABC, and PhosStop inhibitors (Sigma Aldrich) was used to extract the proteins. The lysates for proteomic and PTMomic analysis were heat-inactivated at 90°C for 10 minutes and stored at −80°C until further processing.

### NGS Library preparation for RNA-Seq analysis

The total RNA amount was quantified using the Qubit 2.0 Fluorometric Quantitation system (Thermo Fisher Scientific, Waltham, MA, USA), and the RNA integrity number (RIN) was determined using the Experion Automated Electrophoresis System (Bio-Rad, Hercules, CA, USA). RNA-seq libraries were prepared with the TruSeq Stranded mRNA LT sample preparation kit (Illumina, San Diego, CA, USA) using Sciclone and Zephyr liquid handling workstations (PerkinElmer, Waltham, MA, USA) for pre- and post-PCR steps, respectively. Library concentrations were quantified with the Qubit 2.0 Fluorometric Quantitation system (Life Technologies, Carlsbad, CA, USA). The size distribution was assessed using the Experion Automated Electrophoresis System (Bio-Rad, Hercules, CA, USA). For sequencing, samples were diluted and pooled into NGS libraries in equimolar amounts.

### Next-Generation Sequencing of transcriptome and raw data acquisition

Expression profiling libraries were sequenced on a HiSeq 4000 instrument (Illumina, San Diego, CA, USA) following a 50-base-pair, single-end recipe. Raw data acquisition (HiSeq Control Software, HCS, HD 3.4.0.38) and base calling (Real-Time Analysis Software, RTA, 2.7.7) were performed on the instrument, while the subsequent raw data processing off the instruments involved two custom programs based on Picard tools (2.19.2) (https://github.com/epigen/picard/, https://broadinstitute.github.io/picard/). In the first step, base calls were converted into lane-specific, multiplexed, unaligned BAM files suitable for long-term archival (IlluminaBasecallsToMultiplexSam, 2.19.2-CeMM). In the second step, archive BAM files were demultiplexed into sample-specific, unaligned BAM files (IlluminaSamDemux, 2.19.2-CeMM).

### Transcriptome analysis

NGS reads were mapped to the Genome Reference Consortium GRCh38 assembly via “Spliced Transcripts Alignment to a Reference” (STAR) ^117^ utilizing the “basic” Ensembl transcript annotation from version e100 (April 2020) as reference transcriptome. Since the hg38 assembly flavor of the UCSC Genome Browser was preferred for downstream data processing with Bioconductor packages for entirely technical reasons, Ensembl transcript annotation had to be adjusted to UCSC Genome Browser sequence region names. STAR was run with options recommended by the ENCODE project. Aligned NGS reads overlapping Ensembl transcript features were counted with the Bioconductor (3.11), GenomicAlignments (1.24.0) package via the summarizeOverlaps function in Union mode, taking into account that the Illumina TruSeq stranded mRNA protocol leads to the sequencing of the first strand so that all reads needed inverting before counting. Transcript-level counts were aggregated to gene-level counts, and the Bioconductor DESeq2 (1.28.1) package ^118^ was used to test for differential expression based on a model using the negative binomial distribution.

The initial exploratory analysis included principal component analysis (PCA), multidimensional scaling (MDS), sample distance, and expression heatmap plots, all annotated with variables used in the expression modeling (ggplot2, 3.3.2, and Bioconductor ComplexHeatmap, 2.4.3) ^119^, as well as volcano plots (Bioconductor EnhancedVolcano, 1.6.0). Biologically meaningful results were extracted from the model, log2-fold values were shrunk with the CRAN ashr (2.2.-47) package ^120^, while two-tailed p-values obtained from Wald testing were adjusted with the Bioconductor Independent Hypothesis Weighting (IHW, 1.16.0) package ^121^. The resulting gene lists were annotated and filtered for significantly differentially up- and down-regulated genes.

### Sample preparation for mass spectrometry analysis

The cell lysates were sonicated using a probe sonicator (Branson Digital Sonifier, Branson Ultrasonics Corporation, USA) on ice for 5-10 minutes (20% amplitude, 10 cycles). The lysates were then heated at 95°C and centrifuged at 12,000 rpm each for 10 minutes, respectively. The concentration of proteins was determined by the Bicinchoninic acid (BCA) assay (Pierce, Thermo Fisher Scientific, Waltham, MA). A total of 300 μg proteins per sample were used for downstream processing using the filter-aided sample preparation (FASP) method ^122^. Before FASP, the samples were reduced using Dithiothreitol (DTT) (Sigma Aldrich) at a final concentration of 100 mM at 99°C for 5 minutes, cooled to room temperature, and loaded onto pre-equilibrated VIVACON 500 filter units (Sartorius Stedim Biotech, Germany). Briefly, the lysates were washed with 8M urea in 100 mM Tris/HCl pH 8.5, alkylated with 50 mM Iodoacetamide in 8M Urea, 100 mM Tris/HCl solution, followed by washes with 8M Urea and 50 mM TEAB. The protein samples were finally resuspended in 50 mM TEAB, pH 8.5, and digested with sequencing-grade trypsin (Pierce, Thermo Fisher Scientific) (enzyme to protein ratio 1:50) overnight at 37 °C on a thermomixer. Post digestion, the filter units were centrifuged at 14,000 × g for 20 minutes, followed by 50 mM TEAB and 0.5 M NaCl, respectively, and the process was repeated. Before TMT labeling, the pooled filtrates were acidified with 30% TFA and subjected to solid-phase extraction of peptide digests using Macro Spin columns (Vydac C18, SS18V 30-300μg, The NEST Group, USA).

250 μg SPE purified peptides per sample were labeled with TMT6plex reagent (Thermo Fisher Scientific, USA) following the manufacturer’s instructions. The labeling efficiency and normalization of the mixing ratio were determined following a single injection measurement by LC-MS/MS. The TMT-labeled samples from each channel were pooled to equimolar ratios accordingly. For total proteome analysis, 60 μg of pooled peptides were evaporated to dryness using speedvac and fractionated using SCX StageTip-based fractionation as described earlier ^123^ with minor modifications, including 8 punches of SCX material (3M – 2251) obtained using 14-Gauge needle tip. The remaining pooled TMT labeled peptide digest was subjected to phosphopeptide enrichment.

### Fe-NTA-based phosphopeptides enrichment

Phosphopeptides from each biological replicate were enriched using in-house prepared Fe-NTA microtip columns. Briefly, Ni-NTA Superflow resin (Qiagen, Germany) was activated and coupled with 10 mM Fe(III)chloride solution (Sigma Aldrich, Germany). The activated slurry was resuspended in acetonitrile:methanol: 0.01% acetic acid (1:1:1) solution. Prior to enrichment, the peptides were acidified using 30% TFA and subjected to solid-phase extraction of peptide digests using Macro Spin columns (Vydac C18, SS18V 30-300μg, The NEST Group, USA). The peptides were eluted in SPE-Phospho elution buffer (80% acetonitrile, 0.1% TFA) to achieve a peptide concentration of ~1 μg/μL. The eluate was incubated with an aliquot of activated Fe-NTA slurry on a rotary shaker at room temperature for 60 minutes. This was followed by centrifugation at 6,500 r.p.m. for 1 minute at room temperature and transferring the flow-through to separate tubes for sequential acetyl enrichment. The bound peptides were washed with SPE-Phospho elution buffer thrice. The bound phosphopeptides were released using 50 μL freshly-prepared ammonia solution (~1.4%) and 1.5 μL 100 mM EDTA solution, pH 8, after incubation at room temperature for 10 minutes. After enrichment, the enriched phosphopeptides were loaded onto C8 StageTips to remove contaminating Fe-NTA particles, eluted, and evaporated to dryness using speedvac (Eppendorf, Germany). The enriched phosphopeptides were subjected to offline fractionation using SCX StageTips, as described earlier.

The flow-through from the phosphopeptide enrichment was pooled to 3 fractions, desalted using Waters C18 cartridge, and the eluate was lyophilized. The lyophilized peptide mixtures were dissolved in IAP buffer containing 50 mM MOPS pH 7.2, 10 mM sodium phosphate, and 50 mM NaCl. The Acetyl-lysine motif immunoaffinity beads (Cell Signaling Technology, USA) were washed twice with IAP buffer at 4 °C and then incubated with the peptide mixture for 4 hours with gentle rotation. The unbound peptides were removed by washing the beads with ice-cold IAP buffer (3x) followed by ice-cold water (2x). The enriched peptides were eluted from the beads at room temperature twice using 0.15 % TFA, followed by C18 StageTip-based cleanup before mass spectrometry analysis.

### Liquid chromatography-tandem mass spectrometry analysis

All mass spectrometry data were acquired in centroid mode on a Q Exactive HF Hybrid Quadrupole-Orbitrap mass spectrometer (Thermo Fisher Scientific, Bremen, Germany) coupled to Easy-nLC1200 nano-flow UHPLC (Thermo Scientific, Odense, Denmark). Spray voltage of 1.9 kV was applied with the transfer tube heated to 250°C and S-lens RF of 55%. Internal mass calibration was enabled (lock mass 445.12003 m/z). Tryptic peptides obtained from StageTip-based SCX fractionation from global proteome, phosphoproteome and acetylome fractions were reconstituted in 0.1% formic acid and loaded on an Acclaim PepMap 100 2 cm (3 μm C18 Aq) trap column (Thermo Fisher Scientific). Peptide separation was carried out using Acclaim PepMap 100 C18 HPLC Column, 50 cm heated to 50°C using an integrated column oven. HPLC solvents consisted of 0.1% Formic acid in water (Buffer A) and 0.1% Formic acid, 80% acetonitrile in water (Buffer B). Peptide fractions were eluted at a flow rate of 250nl/min by a non-linear gradient from 6 to 30% B over 110 minutes, followed by a stepwise increase to 60% B in 6 minutes and 95% B in 2 minutes which was held for 20 minutes. Full scan MS spectra (300-1700 m/z) were acquired with a resolution of 120,000 at m/z 200, maximum injection time of 30 ms (total proteome and phosphoproteome), 50 ms (in the case of acetylome), and AGC target value of 3× 10^6^. The 15 most intense precursors with a charge state between 2 and 6 per full scan were selected for fragmentation and isolated with a quadrupole isolation window of 1.2 Th. MS2 analysis was carried out using HCD fragmentation with an NCE of 32% (proteome and acetylome) and 30% (phosphoproteome) and analyzed in the Orbitrap with a resolution of 60,000 at m/z 200, scan range of 200-2000 m/z, AGC target value of 1 x10^5^ and a maximum injection time of 150 ms. Dynamic exclusion was set to 30 secs, and advanced peak determination was deactivated.

### Mass spectrometry data analysis

Mass spectrometry data (.raw) were searched against the human protein database (Uniprot human UP000005640) appended with SARS-CoV-2 protein sequences (Uniprot release 04/09/2020) and common contaminants (245 entries) using SEQUEST HT search algorithm through the Proteome Discoverer platform (v2.4, Thermo Scientific, Bremen, Germany). The search parameter for total proteome included a maximum of two missed cleavages, carbamidomethylation at cysteine, TMT 6-plex Lysine, and TMT 6-plex N-terminal as fixed modifications, oxidation of methionine, Met-loss (Protein N terminus), Acetyl (Protein N terminus), and Met-loss acetyl (Protein N terminus) as dynamic modifications. For phosphoproteome datasets, in addition to the same settings, phosphorylation at serine, threonine, and tyrosine and deamidation at asparagine and glutamine were specified. For acetylome analysis, a maximum of three missed cleavages and acetylation at lysine was selected as a dynamic modification. The precursor mass error tolerance was set at 10 ppm, and the fragment mass error tolerance of 0.05□Da. The data were searched against a decoy database, and a percolator node was employed to calculate the FDR. Peptides identified at a 1% false discovery rate (FDR) were considered further for protein identification. The phosphorylation probabilities at each S/T/Y site and acetylation at K were calculated using the PTM-RS node in the Proteome Discoverer, and peptides with more than 75% site localization probability were considered for further analysis. All peptide groups were normalized for phosphoproteomics and acetylomics datasets by summed intensity normalization and then analyzed on the peptide level. For whole-cell proteomics, normalized PSMs were summed for each accession, and data were exported for further use.

### Metabolomics sample preparation and mass spectrometry analysis

Metabolites were extracted from 100 μL of cell culture medium with 400μL of cold extraction solvent (Acetonitrile:Methanol: Water; 40:40:20). Subsequently, samples were sonicated for 3 cycles with sweep mode (60sec, power 60, and frequency 37), vortexed for 2 minutes, and centrifuged at 4°C, 14,000 rpm for 10 minutes. The supernatant was transferred to autosampler vials for LC-MS analysis. The extracts were analyzed with a Thermo Vanquish UHPLC+ system coupled to a QExactive Orbitrap quadrupole mass spectrometer equipped with a heated electrospray ionization (H-ESI) source probe (M/s Thermo Fischer Scientific, Waltham, MA, USA). A SeQuant ZIC-pHILIC (2.1 × 100 mm, 5-μm particles) HILIC phase analytical column from Merck (M/s Merck KGaA, Darmstadt, Germany) was used as a chromatographic separation column. Gradient elution was carried out with a flow rate of 0.100 mL/minutes using 20 mM ammonium carbonate, adjusted to pH 9.4 with ammonium solution (25%) as mobile phase A and acetonitrile as mobile phase B. The gradient elution was initiated from 20% mobile phase A and 80% of mobile phase B and maintained for 2 minutes. After that, 20% mobile phase A was gradually increased up to 80% till 17 minutes, then mobile phase A was decreased from 80% to 20% in 17.1 minutes and is maintained up to 24 minutes. The column oven and auto-sampler temperatures were set to 40 ± 3 °C and 5 ± 3 °C, respectively. MS equipped with a heated electrospray ionization (H-ESI) source using polarity switching and the following settings: resolution of 35,000, the spray voltages: 4250 V for positive and 3250 V for negative mode, the sheath gas: 25 arbitrary units (AU), and the auxiliary gas: 15 AU, sweep gas flow 0, Capillary temperature: 275°C, S-lens RF level: 50.0. Instrument control was operated with the Xcalibur 4.1.31.9 software (M/s Thermo Fischer Scientific, Waltham, MA, USA). The peaks for metabolite were confirmed using Mass spectrometry metabolite library kit MSMLS-1EA (Ms Sigma Aldrich supplied by IROA Technologies). Peak integration was done with the TraceFinder 4.1 software (M/s Thermo Fischer Scientific, Waltham, MA, USA). The peak area data was exported as an Excel file for further analysis. Data quality was monitored throughout the run using pooled healthy human serum as Quality Control (QC) processed and extracted in the same way as unknown samples and interspersed throughout the run as every 10^th^ sample. After integrating QC data with TraceFinder 4.1, each detected metabolite was checked and %RSD was calculated, and the acceptance limit was set ≤ 20%. Blank samples for carryover were injected after every fifth unknown samples to monitor the metabolites’ carryover effect and calculated against the mean QC area, and the acceptance % carryover limit was set ≤ 20% for each metabolite. Background noise blank (first solvent blank of the run) was injected, and % background noise was calculated against the mean QC area, and the acceptance % background noise limit was set ≤ 20% for each metabolite.

### Cytokine array analysis

The cell culture supernatants from mock-infected and hCoV-19 -S15 strain infected CALU3 cells were collected at indicated time points (0, 3, 6-, 12-, 24-, and 48-hours post-infection (hpi) and subjected to centrifugation for 5 minutes at 14,000 rpm. Cytokines were analyzed using Proteome Profiler Human Cytokine Array Kit (R&D Systems) according to the manufacturer’s instructions. The membranes were exposed to x-ray films, scanned, and analyzed using ImageJ software (NIH). The fold change was calculated in comparison to the mock-infected sample.

### Bioinformatics analysis

The transcriptome FPKM data was subjected to differential analysis. Differentials at each time point were determined using a p-value cut-off of <- 0.01 and log2(fold-change) cut-off of +- 1.5. Protein abundances and phosphosite abundances across multiple replicates were subjected to quantile normalization and differential expression using limma (v3.44.3) in R (v4.0.2 https://www.r-project.org/). Log2 fold changes and p-values were calculated using the proDA (v1.2.0) package for R. Differentials at each time point were determined using a p-value cut-off of <- 0.05and log2(fold-change) cut-off of +- 1.5. For the Acetylome data, abundance ratios obtained from Proteome Discoverer were considered for further analysis. The proteomics and phosphoproteomics datasets generated in the current study were compared with corresponding datasets from Thorne *et al*. ^27^, Hekman *et al*. ^31^, Stukalov *et al*. ^29^ and Grossegesse *et al.^44^* Gene ontology and Pathway analysis for transcriptome total proteome, phosphoproteome and acetylome data were carried out using enrichR (v3.0) R package. The enrichment databases consisted of “GO_Biological_Process_2015”, “GO_Cellular_Component_2015”, and “Reactome_2015” and significant enrichment used a p-value cut-off of 0.05. A list of ISGs was compiled based on previous literature ^27, 124^. The gene sets pertaining to cytokines, chemokines, Hippo signaling, regulation of Hippo signaling, DNA damage response, DNA repair, protein ubiquitination, Regulation of protein mono and polyubiquitination, Alternative splicing by spliceosome, and Regulation of mRNA splicing via spliceosome were obtained from MSigDB (v7.5.1), and comparisons were carried out. The differentials from the phosphoproteome data were subjected to gene set enrichment analysis (GSEA) against the PTM signatures database (PTMsigDB) ^125^, which consists of modification site-specific signatures of perturbations kinase activities, and signaling pathways curated from the literature.

Peak areas for the assayed metabolites were obtained for the specified time points and replicates for the metabolomics data. Fold changes were calculated by dividing the peak area for each time point by the peak area for the unstimulated sample and carrying out the log transformation to the base 2. Replicate values for each time-point were averaged. Metabolites with log2 fold-change values ≥ 1 were considered to be increased, whereas those with log2 fold-change values ≤ −1 were considered to be decreased compared to unstimulated samples. The metabolites increased at each time point and were subjected to metabolite set enrichment analysis (MSEA). Overrepresentation-based MSEA was carried out against the KEGG database using Metaboanalyst 5.0 (https://www.metaboanalyst.ca/)^126^. The individual MSEA results were joined, and the p-value was plotted as a heatmap in R using the pheatmap package (v1.0.12). 2D structures of key metabolites were fetched manually from ChemSpider (http://www.chemspider.com/).

### Kinase motif prediction

Kinase motifs of phosphopeptides from SARS-CoV-2 proteins were predicted using NetPhos 3.1 ^127^ using the SARS-CoV-2 protein fasta file downloaded from Uniprot, which was also used for the proteomics data analysis. Only kinases with a score above 0.5 were considered positive hits.

### Structural analysis of phosphorylation sites

Molecular graphics and analyses were performed with UCSF ChimeraX ^128^. Experimental models were used for ORF7a (PDB ID 6w37) (doi: 10.2210/pdb6w37/pdb), ORF8 (PDB ID 7jx6) (doi: 10.2210/pdb7jx6/pdb), ORF9b (PDB ID 7kdt) ^129^. Models for full-length ORF3a, ORF6, M, N, Rep1a residues 1-180 (Nsp1) and residues 2483-2667 (Nsp3) were obtained using TrRosetta through the Robetta server. For visualization, regions of Robetta models present in the experimental structures of ORF3a (PDB ID 7kjr) ^130^, N NTD (PDB ID 6vyo) (10.2210/pdb6VYO/pdb) and CTD (PDB ID 6wzo) ^84^, Rep1a Nsp1 NTD (PDB ID 7k7p) ^131^ and CTD (PDB ID 7jqb)^132^ were hidden in the illustrations and replaced by the experimental models. Structural models of SARS-CoV-2 proteins were visualized as cartoon representations and surfaces were colored by electrostatic potential.

### Western blot analysis

Protein samples were run on pre-cast NuPAGE™ Bis-Tris gels (Invitrogen) with 1x MOPS buffer (Invitrogen) and transferred on nitrocellulose membranes using the iBlot®2 Gel Transfer Device (Invitrogen). Membranes were washed in Tris Buffered Saline with 0.1% Tween-X100 (TBS-T) and blocked with TBS-T containing 5% bovine serum albumin (BSA, Sigma-Aldrich). Membranes were incubated with primary antibodies at 4°C overnight. The following primary antibodies were used:, anti-ß-Actin (1:5000; cat#6276; Abcam), anti-OAS1 (1:1000; cat#14498; Cell Signaling Technology), anti-ISG15 (1:1000; cat#2734S; Cell Signaling Technology), anti-TRIM5α (1:1000; cat#14326; Cell Signaling Technology), anti-RNase-L (1:1000; cat#27281; Cell Signaling Technology), anti-MX1 (1:1000; cat#37849; Cell Signaling Technology), anti-TYKi (CMPK2) (1:1000; cat#NBP1-80653; Novus Biologicals), anti-p53 (1:1000; cat#; Cell Signaling Technology), anti-phospho p53 (Ser15) (1:1000; cat#; Cell Signaling Technology), anti-phospho γH2AX (Ser139) (1:1000; cat#2577; Cell Signaling Technology), anti-phospho TBK1 (Ser172) (1:1000; cat#5483; Cell Signaling Technology), anti-phospho IRF3 (Ser396) (1:1000; cat#29047; Cell Signaling Technology), anti-phospho JNK (Ser) (1:1000; cat#4668; Cell Signaling Technology), anti-phospho ERK1/2 (Thr202/Tyr204) (1:1000; cat#4370; Cell Signaling Technology), anti-phospho p38 (Thr180/Tyr182) (1:1000; cat#4511; Cell Signaling Technology), anti-phospho STAT1 (Ser727) (1:1000; cat#8826; Cell Signaling Technology), anti-phospho STAT3 (Tyr705) (1:1000; cat#9145S; Cell Signaling Technology), anti-phospho EIF2α (Ser51) (1:1000; cat#3398; Cell Signaling Technology) and anti-phospho YAP1 (Ser61) (1:1000; kind gift from Dr. Wenqi Wang, University of California, Irvine). Membranes were washed in TBS-T and incubated with secondary antibodies (HRP-conjugated, DAKO) for 1 hour at room temperature in TBS-T containing 5% milk or BSA. The blots were developed with SuperSignal West Femto Substrate (Thermo Scientific) and captured with LI-COR Odyssey system (LI-COR Biosciences, Lincoln, NE, USA).

## Data Availability

The RNA-Seq data from the current study are available from the ArrayExpress database under the accession number E-MTAB-12134.

The mass spectrometry proteomics raw data have been deposited to the ProteomeXchange Consortium via the PRIDE partner ^133^ repositories with the dataset identifier PXD032677. Reviewer account details:

Username

Password: CJxc4Mex

The processed data is also available for querying at www.sarscov2map.org

## Acknowledgments

We thank the Proteomics and Modomics Experimental Core Facility (PROMEC), Norwegian University of Science and Technology (NTNU), for support and expertise on LC-MS instrumentation. PROMEC is funded by the Faculty of Medicine and Health Sciences at NTNU and the Central Norway Regional Health Authority. Data storage and handling are supported under the NIRD/Notur project NN9036K. We thank Barbara Van Loon, IKOM, NTNU, for providing us with antibodies to assess cell cycle and DNA damage response. We thank Wenqi Wang, University of California, Irvine, USA for providing pYAP (S61) antibody. This research was funded by the Research Council of Norway (FRIMEDBIO “Young Research Talent” Grant 263168 to R.K.K.; and Centres of Excellence Funding Scheme Project 223255/F50 to CEMIR), Onsager fellowship from NTNU (to R.K.K.). The FIMM Metabolomics Unit is supported by HiLIFE and Biocenter, Finland.

## Author Contributions

Conceptualization, SMP, DK, and RKK.; Methodology, SMP., DK and RKK.; Formal Analysis, SMP, YS, MWG, and RKK.; Investigation, SMP, HK, AN, LH, and DK; Data Curation, YS and MG; Writing – Original Draft, SMP, and YS.; Writing – Review & Editing, SMP, YS, and RKK; Supervision: RKK.; Project Administration: RKK; Funding Acquisition, TE, DK, MB, and RKK.

## Abbreviations

AGC: Automatic Gain Control
ATCC: American Type Culture Collection
BCA: Bicinchoninic acid
BSA: Bovine Serum Albumin
CRISPR: Clustered Regularly Interspaced Short Palindromic Repeats
DEG: Differentially Expressed Genes
DMEM: Dulbecco’s Modified Eagle Medium
DTT: Dithiothreitol
FASP: Filter-Aided Sample Preparation
FBS: Fetal Bovine Serum
FDA: Food and Drug Administration
FDR: False Discovery Rate
HCD: Higher Energy Collision Dissociation
H-ESI: Heated Electrospray ionization
HILIC: Hydrophilic interaction liquid chromatography
hpi: hours post-infection
IAA: Iodoacetamide
ISG: Interferon Stimulated Genes
KEGG: Kyoto Encyclopedia of Genes and Genomes
LC-MS/MS: Liquid Chromatography Tandem Mass Spectrometry
MDS: Multi-Dimensional Scaling
MOI: Multiplicity of Infection
MOPS: 3-(N-morpholino)propanesulfonic acid
mRNA: messenger Ribo Nucleic Acid
MSEA: Metabolite set enrichment analysis
NCE: Normalized collision energy
NGS: Next-Generation Sequencing
PCA: Principal component analysis
PDB: Protein Data Bank
PPI: Protein-protein interaction
PSM: Peptide-spectrum match
PTM: Post-Translational Modification
QC: Quality Control
RSD: Relative standard deviation
SARS-CoV-2: Severe Acute Respiratory Syndrome Coronavirus 2
SCX: Strong cation exchange
SDS: Sodium dodecyl-sulfate
SPE: Solid-Phase Extraction
TBS: Tris-Buffered Saline
TEAB: Triethylammonium bicarbonate
TFA: Trifluoroacetic acid
TMT: Tandem mass tag
UHPLC: Ultra-high performance liquid chromatography

