## Supplementary Figure for "MultiOMICs landscape of SARS-CoV-2-induced host responses in human lung epithelial cells"

**Supplementary Data**

**Supplementary Tables**

**Supplementary Table S1.** List of differentially regulated transcripts identified in Calu-3 cells infected with SARS-CoV-2/ Trondheim-S15/2020 strain at time intervals 3, 6, 12, 24, and 48 hpi.

**Supplementary Table S2.** List of differentially expressed proteins identified in Calu-3 cells infected with SARS-CoV-2/ Trondheim-S15/2020 strain at 3, 6, 12, 24, and 48 (hpi).

**Supplementary Table S3.** List of differentially phosphorylated proteins and phosphosites identified in Calu-3 cells infected with SARS-CoV-2/ Trondheim-S15/2020 strain at 3, 6, 12, 24, and 48 (hpi).

**Supplementary Table S4.** List of differentially acetylated proteins and acetylation sites identified in Calu-3 cells infected with SARS-CoV-2/ Trondheim-S15/2020 strain at 3, 6, 12, 24, and 48 (hpi).

**Supplementary Table S5.** Pathways enriched in the 8 phosphoproteome clusters

**Supplementary Table S6.** Exometabolomics data from Calu-3 cells infected with SARS-CoV-2 3, 6, 12, 24, and 48 (hpi).

**Supplementary Table S7.** Viral omics profile from Calu-3 cells infected with SARS-CoV-2 at 3, 6, 12, 24, and 48 (hpi).

**Supplementary Figures**

**Supplementary Figure S1.** Correlation graphs depicting replicate samples and various time points for SARS-CoV-2/ Trondheim-S15/2020 strain infected Calu-3 cell **A.** transcriptome **B.** Total Proteome, **C.** Phosphoproteome, and **D.** acetylome data.

**Supplementary Figure S2. A.** Biological Processes (GO-BP) enriched in the Calu-3 transcriptome and proteome after SARS-CoV-2 infection. **B.** An upset graph illustrating overlap of differentially expressed proteins in lung cell lines in response to SARS-CoV-2 infection between the current study and various published datasets, including Thorn *et al.*, Hekman *et al.*, Grossegesse *et al.*, and Stukalov *et al.* **C.** Heatmap depicting transcript levels of cytokines and chemokines in Calu-3 cells at time intervals 3, 6, 12, 24, and 48 hpi. **D.** A scan of the cytokine array showing changes in levels of cytokines in Calu-3 cells infected with SARS-CoV-2/ Trondheim-S15/2020 strain.

**Supplementary Figure S3. A.** An upset graph showing overlap of hyperphosphorylated proteins in lung cell lines in response to SARS-CoV-2 infection between the current study and various published datasets, including Thorn *et al.*, Hekman *et al.*, and Stukalov *et al.* **B.**

Heatmap showing enriched upstream kinases from the phosphoproteomics profile of Calu-3 cells upon SARS-CoV-2 infection **C.** Statistics of differentially phosphorylated transcription factors in response to SARS-CoV-2 infection **D.** Heatmap showing changes in the levels of transcription factors after SARS-CoV-2 infection. **E.** Graph showing changes in Lysine (K) acetylation sites, Serine (S) phosphorylation sites and protein abundance of Vimentin (VIM). **F.** Graph showing changes in Lysine (K) acetylation and ubiquitination sites of leucine rich repeat containing 59 (LRRC59)

**Supplementary Figure S4.** Metabolite set enrichment for Calu-3 exometabolites changing after infection with SARS-CoV-2 infection at **A.** 3, **B.** 6, **C.** 12, **D.** 24, and **E.** 48 hpi.

**Supplementary Figure S5.** Phosphorylation sites unique to this study are presented as magenta sticks, whereas common sites are in cyan. Experimental structural models of **A.** ORF7a (PDB ID 6w37), **B.** ORF8 (PDB ID 7jx6), **C.** ORF9b (PDB ID 7kdt) and Robetta models of **D.** N protein **E.** ORF3a, **F.** ORF6, **G.** Rep1a, **H.** Nsp3, and **I.** Protein M are shown in rainbow cartoon representation (N-to-C blue to red). Missing regions of Protein N and Rep1a Nsp1 were modelled with Robetta and presented as a rainbow cartoon and were hidden where experimental structures of Protein N (PDB ID 6vyo for NTD, in olive and PDB ID 6wzo for CTD, in grey) and Rep1a Nsp1 (PDB ID 7k7p for NTD, in teal and PDB ID 7jqb for CTD, in purple) were overlapping with the models. The second protomer of ORF3a (PDB ID 7kjr in grey) and Tom70, the interaction partner of ORF9b (PDB ID 7kdt, in tan) are shown in uniform colors. Selected hydrogen bonds are shown as cyan dotted lines. The figure was prepared with ChimeraX (Goddard et al doi: 10.1002/pro.3235)

**Supplementary Figure S6.** Graphs displaying the average trend of differentials from transcriptomics, proteomics, and phosphoproteomics datasets with respect to various pathways and processes, including **A.** Alternative splicing by spliceosome **B.** Regulation of mRNA splicing via spliceosome. Heatmaps showing differential changes of transcripts, proteins and phosphorylation sites for **C.** Hippo signaling **D.** Regulation of Hippo signaling, **E.** DNA damage response, **F.** DNA repair, **G.** Protein ubiquitination, **H.** Regulation of protein mono and polyubiquitination **I.** Graph showing protein expression levels of Ubiquitin (Ub) E2 and E3 ligases in response to SARS-CoV-2 infection. **J.** Heatmap showing changes in proteins involved in regulation of the cell cycle.

Figure S1

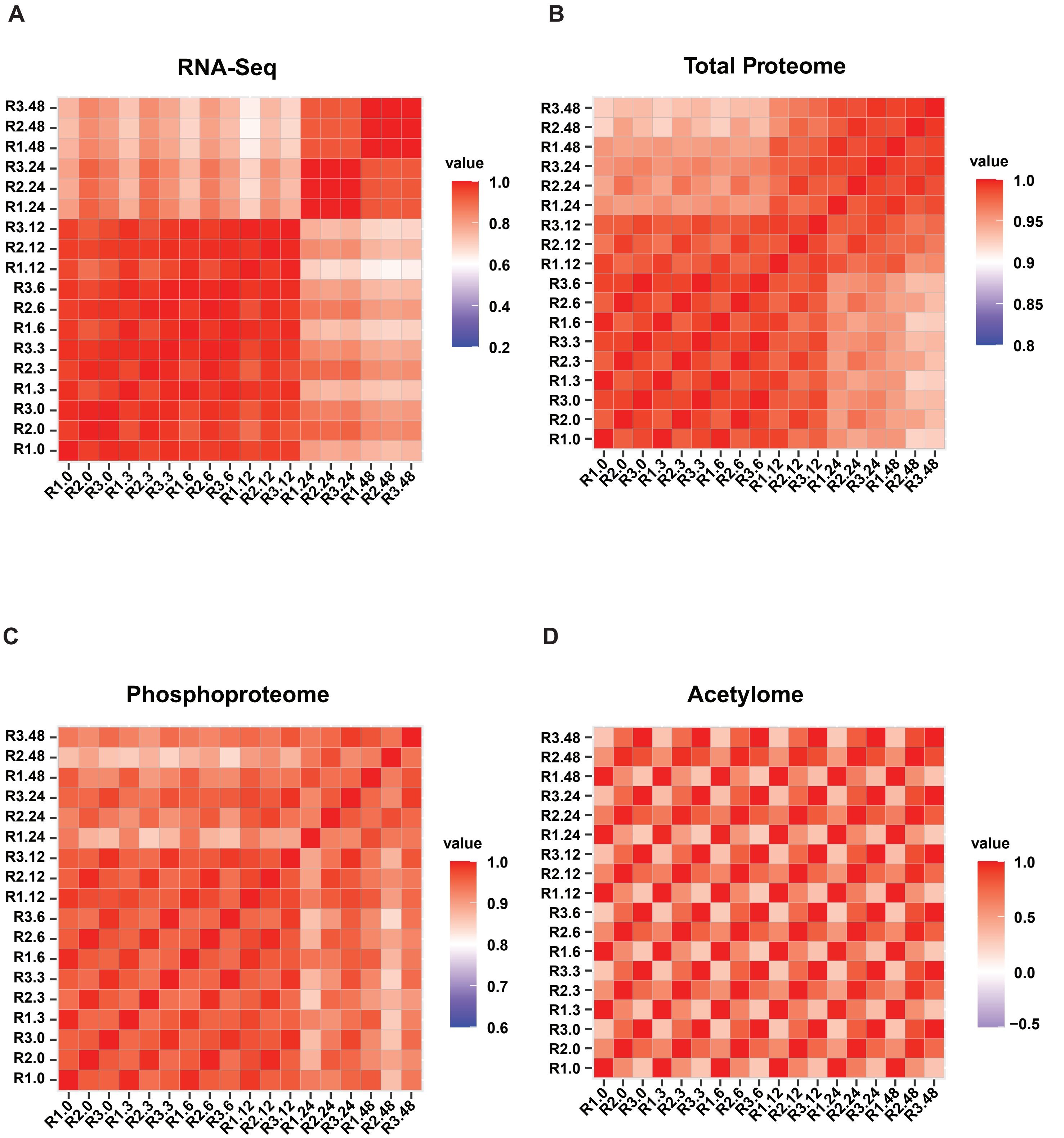
A
RNA-Seq
R3.48
R2.48
R1.48
R3.24
R2.24
R1.24
R3.12
R2.12
R1.12
R3.6
R2.6
R1.6
R3.3
R2.3
R1.3
R3.0
R2.0
R1.0
R1.0 R2.0 R3.0 R1.3 R2.3 R3.3 R1.6 R2.6 R3.6 R1.12 R2.12 R3.12 R1.24 R2.24 R3.24 R1.48 R2.48 R3.48
value
1.0
0.8
0.6
0.4
0.2
B
Total Proteome
value
1.0
0.95
0.9
0.85
0.8
C
Phosphoproteome
value
1.0
0.9
0.8
0.7
0.6
D
Acetylome
value
1.0
0.5
0.0
−0.5

**Figure S2**

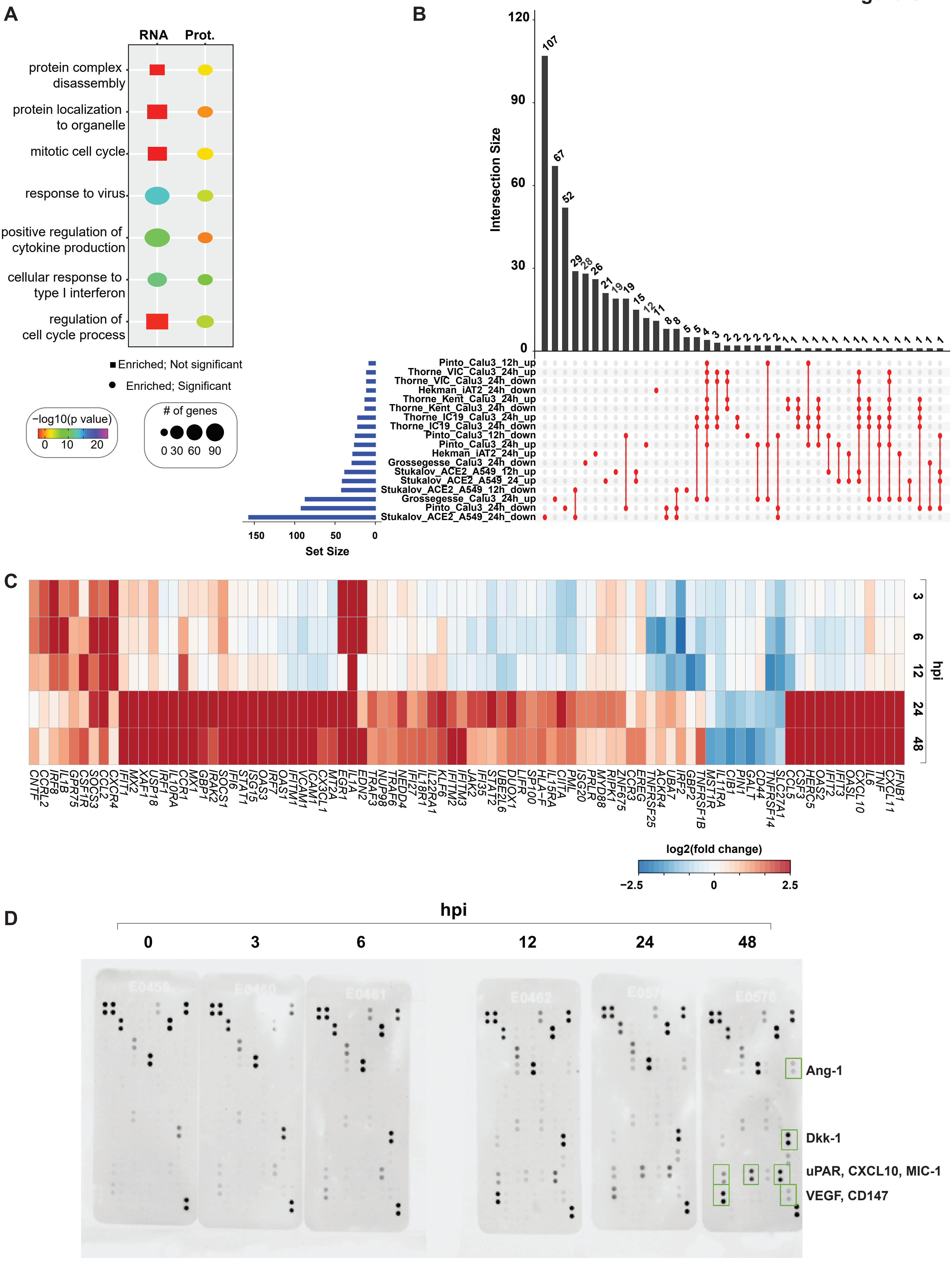

A
RNA
Prot.
protein complex disassembly
protein localization to organelle
mitotic cell cycle
response to virus
positive regulation of cytokine production
cellular response to type I interferon
regulation of cell cycle process
Enriched; Not significant
Enriched; Significant
−log10(p value)
0 10 20
### of genes
0 30 60 90
B
Intersection Size
120
90
60
30
0
107 67 52 29 28 26 21 19 19 15 12 11 8 8 5 5 4 3 2 2 2 2 2 2 1 1 1 1 1 1 1 1 1 1 1 1 1 1 1 1
Pinto_Calu3_12h_up
Thorne_VIC_Calu3_24h_up
Thorne_VIC_Calu3_24h_down
Hekman_iAT2_24h_down
Thorne_Kent_Calu3_24h_up
Thorne_Kent_Calu3_24h_down
Thorne_IC19_Calu3_24h_up
Thorne_IC19_Calu3_24h_down
Pinto_Calu3_12h_down
Pinto_Calu3_24h_up
Hekman_iAT2_24h_up
Grossegesse_Calu3_24h_down
Stukalov_ACE2_A549_12h_up
Stukalov_ACE2_A549_24_up
Stukalov_ACE2_A549_12h_down
Grossegesse_Calu3_24h_up
Pinto_Calu3_24h_down
Stukalov_ACE2_A549_24h_down
150 100 50 0
Set Size
C
3
6
12
24
48
hpi
CNTF CCRL2 IRF8 IL1B GPR75 CSF1R SOCS3 CCL2 CXCR4 IFIT1 MX2 XAF1 USP18 IRF1 IL10RA CCR1 MX1 GBP1 IRAK2 SOCS1 IFI6 STAT1 ISG15 OAS3 IRF7 OAS1 IFITM1 VCAM1 ICAM1 CX3CL1 MT2A EGR1 IL1A EDN2 TRAF3 NUP98 TRAF6 NEDD4 IFI27 IL18R1 IL22RA1 KLF6 IFITM2 IFITM3 JAK2 IFI35 STAT2 UBE2L6 DUOX1 LIFR SP100 HLA-F IL15RA CIITA PML ISG20 PRLR MYD88 RIPK1 ZNF675 CCR3 EREG TNFRSF25 ACKR4 UBA7 IRF2 GBP2 TNFRSF1B MST1R IL11RA CIB1 PIN1 GALT CD44 TNFRSF14 SLC27A1 CCL5 CSF3 HERC5 OAS2 IFIT2 IFIT3 OASL CXCL10 IL6 TNF CXCL11 IFNB1
log2(fold change)
−2.5 0 2.5
D
hpi
0 3 6 12 24 48
Ang-1
Dkk-1
uPAR, CXCL10, MIC-1
VEGF, CD147

**Figure S3**

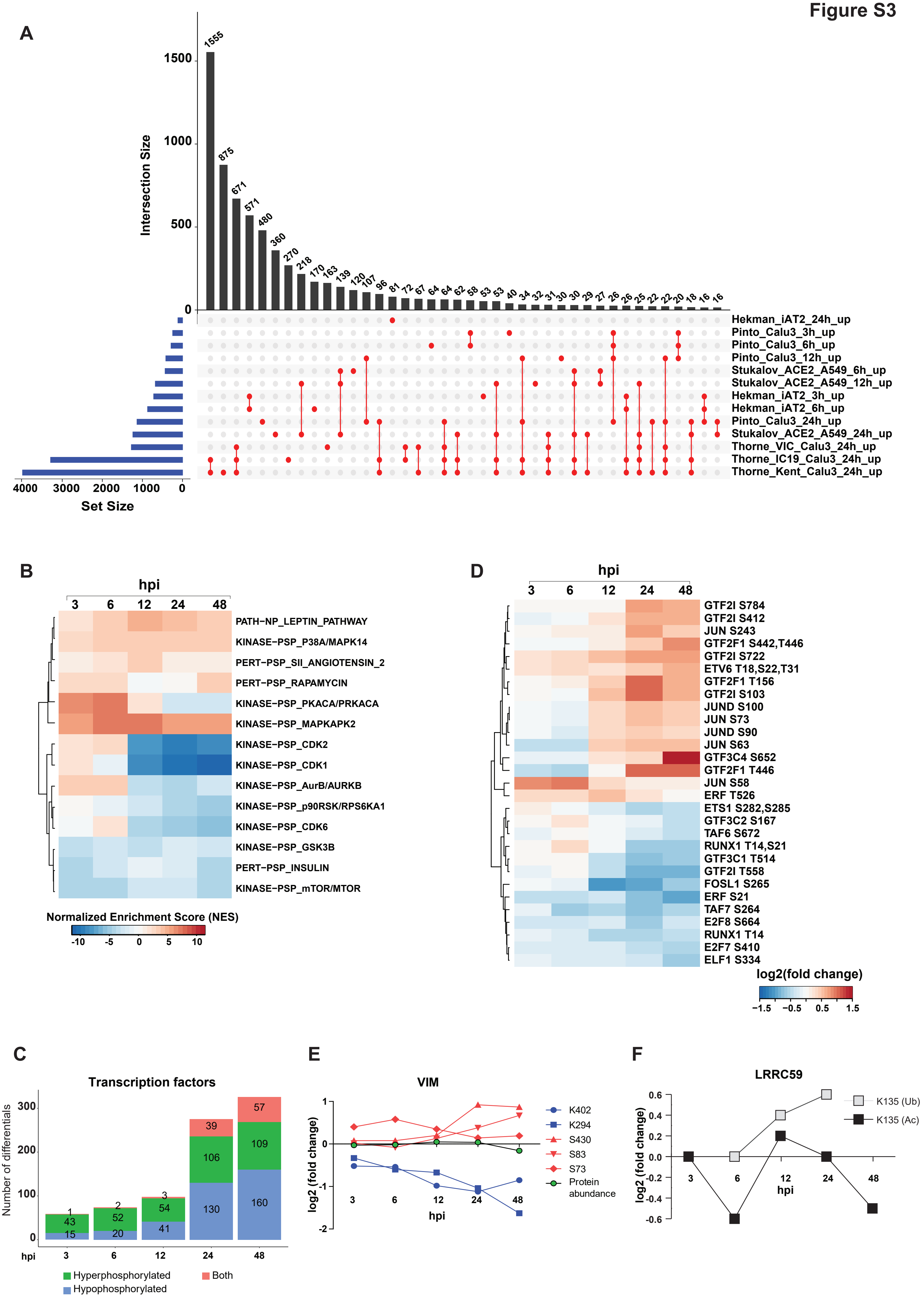
A
Intersection Size
1555
875
671
571
480
360
270
218
170
163
139
120
107
96
81
72
67
64
64
62
58
53
53
40
34
32
31
30
30
29
27
26
26
25
22
22
20
18
16
16
0
500
1000
1500
Hekman_iAT2_24h_up
Pinto_Calu3_3h_up
Pinto_Calu3_6h_up
Pinto_Calu3_12h_up
Stukalov_ACE2_A549_6h_up
Stukalov_ACE2_A549_12h_up
Hekman_iAT2_3h_up
Hekman_iAT2_6h_up
Pinto_Calu3_24h_up
Stukalov_ACE2_A549_24h_up
Thorne_VIC_Calu3_24h_up
Thorne_IC19_Calu3_24h_up
Thorne_Kent_Calu3_24h_up
4000
3000
2000
1000
0
Set Size
B
hpi
3
6
12
24
48
PATH−NP_LEPTIN_PATHWAY
KINASE−PSP_P38A/MAPK14
PERT−PSP_SII_ANGIOTENSIN_2
PERT−PSP_RAPAMYCIN
KINASE−PSP_PKACA/PRKACA
KINASE−PSP_MAPKAPK2
KINASE−PSP_CDK2
KINASE−PSP_CDK1
KINASE−PSP_AurB/AURKB
KINASE−PSP_p90RSK/RPS6KA1
KINASE−PSP_CDK6
KINASE−PSP_GSK3B
PERT−PSP_INSULIN
KINASE−PSP_mTOR/MTOR
Normalized Enrichment Score (NES)
-10
-5
0
5
10
D
hpi
3
6
12
24
48
GTF2I S784
GTF2I S412
JUN S243
GTF2F1 S442,T446
GTF2I S722
ETV6 T18,S22,T31
GTF2F1 T156
GTF2I S103
JUND S100
JUN S73
JUND S90
JUN S63
GTF3C4 S652
GTF2F1 T446
JUN S58
ERF T526
ETS1 S282,S285
GTF3C2 S167
TAF6 S672
RUNX1 T14,S21
GTF3C1 T514
GTF2I T558
FOSL1 S265
ERF S21
TAF7 S264
E2F8 S664
RUNX1 T14
E2F7 S410
ELF1 S334
log2(fold change)
−1.5
−0.5
0.5
1.5
C
Transcription factors
Number of differentials
300
200
100
0
1
43
15
2
52
20
3
54
41
39
106
130
57
109
160
hpi
3
6
12
24
48
Hyperphosphorylated
Hypophosphorylated
Both
E
VIM
log2 (fold change)
1
0
-1
-2
3
6
12
24
48
hpi
K402
K294
S430
S83
S73
Protein abundance
F
LRRC59
log2 (fold change)
0.6
0.4
0.2
0.0
-0.2
-0.4
-0.6
3
6
12
24
48
hpi
K135 (Ub)
K135 (Ac)

**Figure S4**

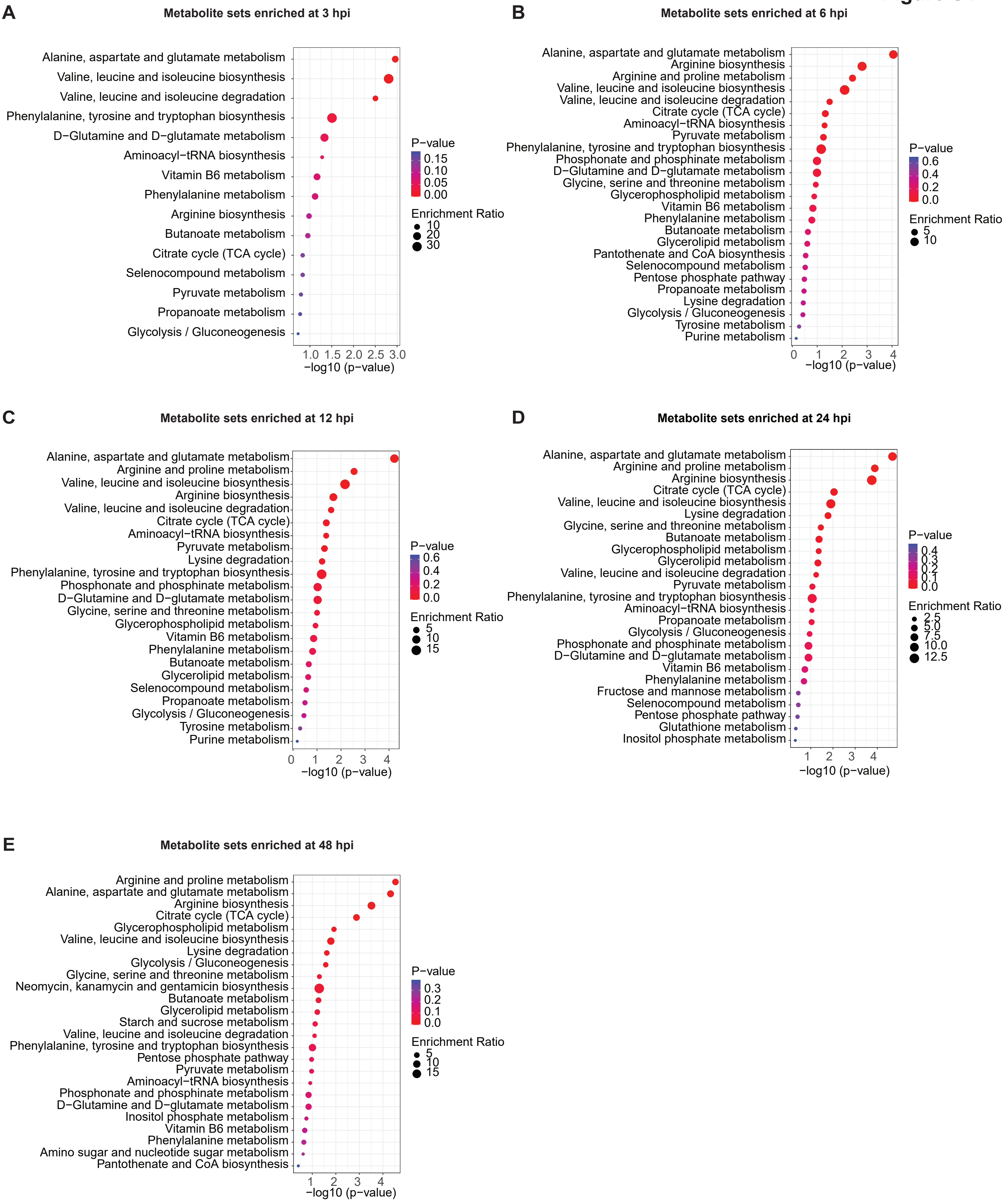
A
Metabolite sets enriched at 3 hpi
Alanine, aspartate and glutamate metabolism
Valine, leucine and isoleucine biosynthesis
Valine, leucine and isoleucine degradation
Phenylalanine, tyrosine and tryptophan biosynthesis
D−Glutamine and D−glutamate metabolism
Aminoacyl−tRNA biosynthesis
Vitamin B6 metabolism
Phenylalanine metabolism
Arginine biosynthesis
Butanoate metabolism
Citrate cycle (TCA cycle)
Selenocompound metabolism
Pyruvate metabolism
Propanoate metabolism
Glycolysis / Gluconeogenesis
1.0 1.5 2.0 2.5 3.0
−log10 (p−value)
P−value
0.15
0.10
0.05
0.00
Enrichment Ratio
10
20
30
B
Metabolite sets enriched at 6 hpi
Alanine, aspartate and glutamate metabolism
Arginine biosynthesis
Arginine and proline metabolism
Valine, leucine and isoleucine biosynthesis
Valine, leucine and isoleucine degradation
Citrate cycle (TCA cycle)
Aminoacyl−tRNA biosynthesis
Pyruvate metabolism
Phenylalanine, tyrosine and tryptophan biosynthesis
Phosphonate and phosphinate metabolism
D−Glutamine and D−glutamate metabolism
Glycine, serine and threonine metabolism
Glycerophospholipid metabolism
Vitamin B6 metabolism
Phenylalanine metabolism
Butanoate metabolism
Glycerolipid metabolism
Pantothenate and CoA biosynthesis
Selenocompound metabolism
Pentose phosphate pathway
Propanoate metabolism
Lysine degradation
Glycolysis / Gluconeogenesis
Tyrosine metabolism
Purine metabolism
0 1 2 3 4
−log10 (p−value)
P−value
0.6
0.4
0.2
0.0
Enrichment Ratio
5
10
C
Metabolite sets enriched at 12 hpi
Alanine, aspartate and glutamate metabolism
Arginine and proline metabolism
Valine, leucine and isoleucine biosynthesis
Arginine biosynthesis
Valine, leucine and isoleucine degradation
Citrate cycle (TCA cycle)
Aminoacyl−tRNA biosynthesis
Pyruvate metabolism
Lysine degradation
Phenylalanine, tyrosine and tryptophan biosynthesis
Phosphonate and phosphinate metabolism
D−Glutamine and D−glutamate metabolism
Glycine, serine and threonine metabolism
Glycerophospholipid metabolism
Vitamin B6 metabolism
Phenylalanine metabolism
Butanoate metabolism
Glycerolipid metabolism
Selenocompound metabolism
Propanoate metabolism
Glycolysis / Gluconeogenesis
Tyrosine metabolism
Purine metabolism
0 1 2 3 4
−log10 (p−value)
P−value
0.6
0.4
0.2
0.0
Enrichment Ratio
5
10
15
D
Metabolite sets enriched at 24 hpi
Alanine, aspartate and glutamate metabolism
Arginine and proline metabolism
Arginine biosynthesis
Citrate cycle (TCA cycle)
Valine, leucine and isoleucine biosynthesis
Lysine degradation
Glycine, serine and threonine metabolism
Butanoate metabolism
Glycerophospholipid metabolism
Glycerolipid metabolism
Valine, leucine and isoleucine degradation
Pyruvate metabolism
Phenylalanine, tyrosine and tryptophan biosynthesis
Aminoacyl−tRNA biosynthesis
Propanoate metabolism
Glycolysis / Gluconeogenesis
Phosphonate and phosphinate metabolism
D−Glutamine and D−glutamate metabolism
Vitamin B6 metabolism
Phenylalanine metabolism
Fructose and mannose metabolism
Selenocompound metabolism
Pentose phosphate pathway
Glutathione metabolism
Inositol phosphate metabolism
1 2 3 4
−log10 (p−value)
P−value
0.4
0.3
0.2
0.1
0.0
Enrichment Ratio
2.5
5.0
7.5
10.0
12.5
E
Metabolite sets enriched at 48 hpi
Arginine and proline metabolism
Alanine, aspartate and glutamate metabolism
Arginine biosynthesis
Citrate cycle (TCA cycle)
Glycerophospholipid metabolism
Valine, leucine and isoleucine biosynthesis
Lysine degradation
Glycolysis / Gluconeogenesis
Glycine, serine and threonine metabolism
Neomycin, kanamycin and gentamicin biosynthesis
Butanoate metabolism
Glycerolipid metabolism
Starch and sucrose metabolism
Valine, leucine and isoleucine degradation
Phenylalanine, tyrosine and tryptophan biosynthesis
Pentose phosphate pathway
Pyruvate metabolism
Aminoacyl−tRNA biosynthesis
Phosphonate and phosphinate metabolism
D−Glutamine and D−glutamate metabolism
Inositol phosphate metabolism
Vitamin B6 metabolism
Phenylalanine metabolism
Amino sugar and nucleotide sugar metabolism
Pantothenate and CoA biosynthesis
1 2 3 4
−log10 (p−value)
P−value
0.3
0.2
0.1
0.0
Enrichment Ratio
5
10
15

**Figure S5**

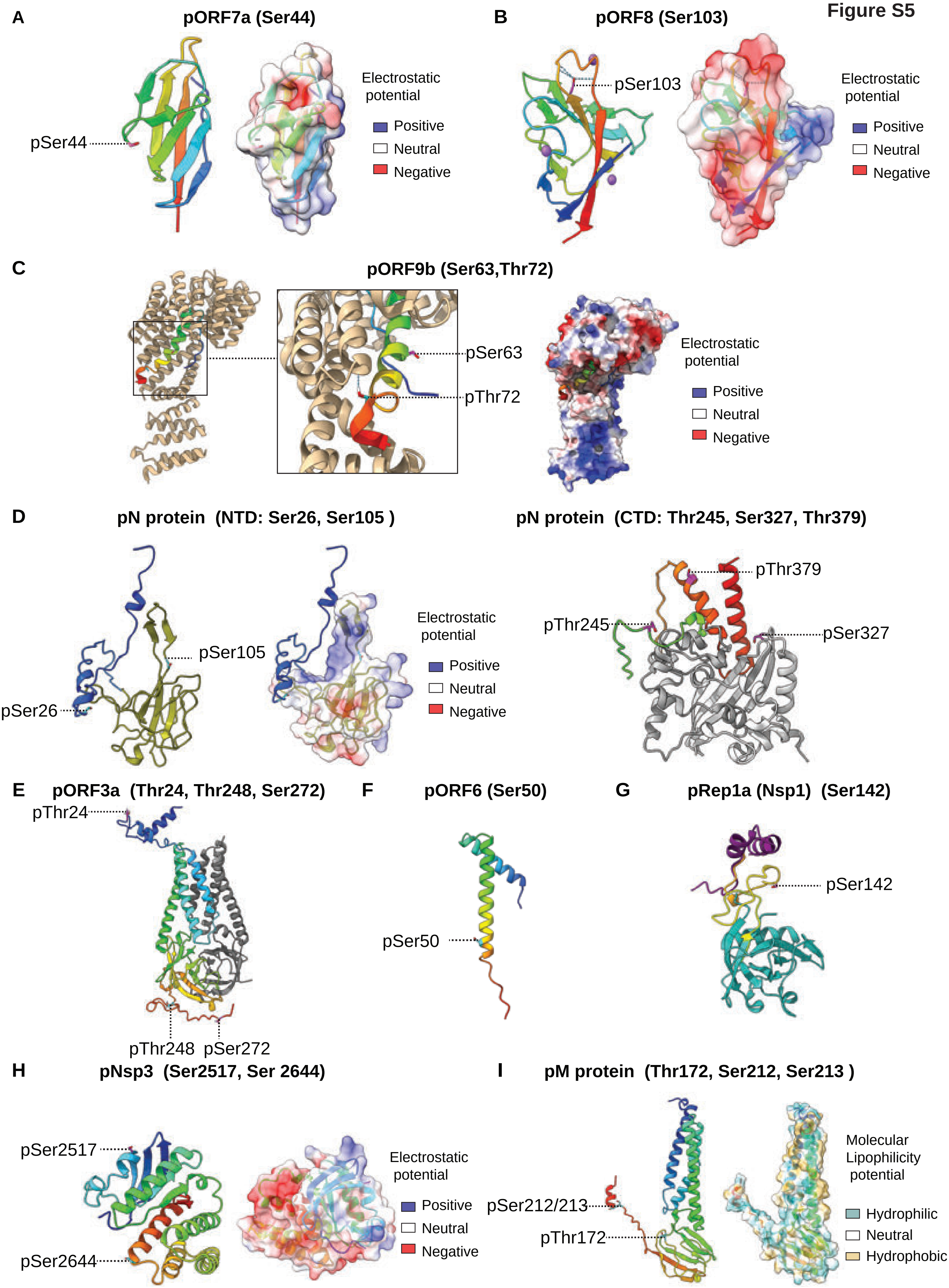

A
pORF7a (Ser44)
pSer44
Electrostatic potential
Positive
Neutral
Negative
B
pORF8 (Ser103)
pSer103
Electrostatic potential
Positive
Neutral
Negative
C
pORF9b (Ser63,Thr72)
pSer63
pThr72
Electrostatic potential
Positive
Neutral
Negative
D
pN protein (NTD: Ser26, Ser105 )
pSer105
pSer26
Electrostatic potential
Positive
Neutral
Negative
pN protein (CTD: Thr245, Ser327, Thr379)
pThr379
pThr245
pSer327
E
pORF3a (Thr24, Thr248, Ser272)
pThr24
pThr248
pSer272
F
pORF6 (Ser50)
pSer50
G
pRep1a (Nsp1) (Ser142)
pSer142
H
pNsp3 (Ser2517, Ser 2644)
pSer2517
pSer2644
Electrostatic potential
Positive
Neutral
Negative
I
pM protein (Thr172, Ser212, Ser213 )
pSer212/213
pThr172
Molecular Lipophilicity potential
Hydrophilic
Neutral
Hydrophobic

Figure S6

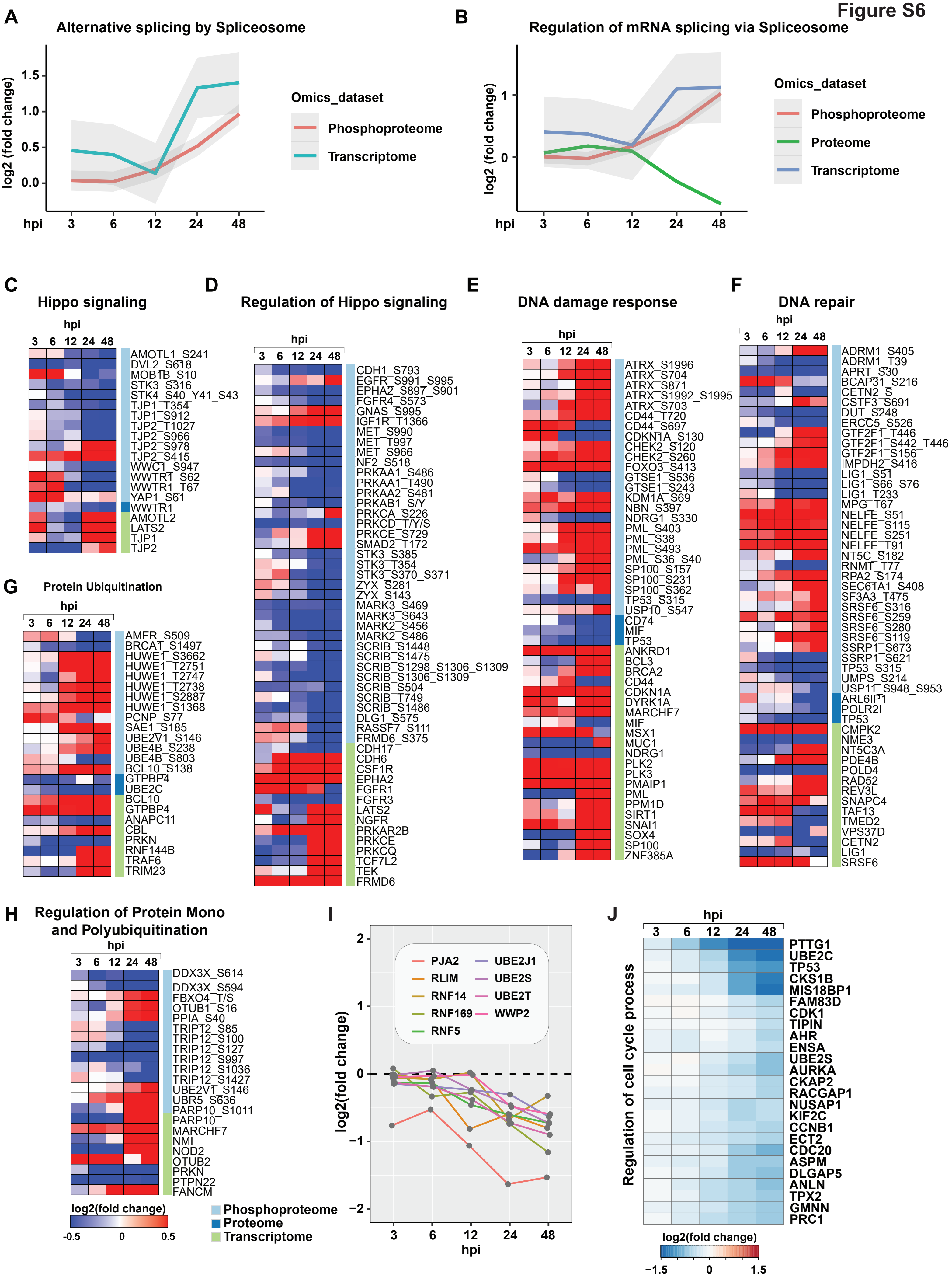
A
Alternative splicing by Spliceosome
log2 (fold change)
1.5
1.0
0.5
0.0
hpi
3
6
12
24
48
Omics_dataset
Phosphoproteome
Transcriptome
B
Regulation of mRNA splicing via Spliceosome
log2 (fold change)
1
0
hpi
3
6
12
24
48
Omics_dataset
Phosphoproteome
Proteome
Transcriptome
C
Hippo signaling
hpi
3 6 12 24 48
AMOTL1_S241
DVL2_S618
MOB1B_S10
STK3_S316
STK4_S40_Y41_S43
TJP1_T354
TJP1_S912
TJP2_T1027
TJP2_S966
TJP2_S978
TJP2_S415
WWC1_S947
WWTR1_S62
WWTR1_T67
YAP1_S61
WWTR1
AMOTL2
LATS2
TJP1
TJP2
D
Regulation of Hippo signaling
hpi
3 6 12 24 48
CDH1_S793
EGFR_S991_S995
EPHA2_S897_S901
FGFR4_S573
GNAS_S995
IGF1R_T1366
MET_S990
MET_T997
MET_S966
NF2_S518
PRKAA1_S486
PRKAA1_T490
PRKAA2_S481
PRKAB1_S/Y
PRKCA_S226
PRKCD_T/Y/S
PRKCE_S729
SMAD2_T172
STK3_S385
STK3_T354
STK3_S370_S371
ZYX_S281
ZYX_S143
MARK3_S469
MARK3_S643
MARK2_S456
MARK2_S486
SCRIB_S1448
SCRIB_S1475
SCRIB_S1298_S1306_S1309
SCRIB_S1306_S1309
SCRIB_S504
SCRIB_T749
SCRIB_S1486
DLG1_S575
RASSF7_S111
FRMD6_S375
CDH17
CDH6
CSF1R
EPHA2
FGFR1
FGFR3
LATS2
NGFR
PRKAR2B
PRKCE
PRKCQ
TCF7L2
TEK
FRMD6
E
DNA damage response
hpi
3 6 12 24 48
ATRX_S1996
ATRX_S704
ATRX_S871
ATRX_S1992_S1995
ATRX_S703
CD44_T720
CD44_S697
CDKN1A_S130
CHEK2_S120
CHEK2_S260
FOXO3_S413
GTSE1_S536
GTSE1_S243
KDM1A_S69
NBN_S397
NDRG1_S330
PML_S403
PML_S38
PML_S493
PML_S36_S40
SP100_S157
SP100_S231
SP100_S362
TP53_S315
USP10_S547
CD74
MIF
TP53
ANKRD1
BCL3
BRCA2
CD44
CDKN1A
DYRK1A
MARCHF7
MIF
MSX1
MUC1
NDRG1
PLK2
PLK3
PMAIP1
PML
PPM1D
SIRT1
SNAI1
SOX4
SP100
ZNF385A
F
DNA repair
hpi
3 6 12 24 48
ADRM1_S405
ADRM1_T39
APRT_S30
BCAP31_S216
CETN2_S
CSTF3_S691
DUT_S248
ERCC5_S526
GTF2F1_T446
GTF2F1_S442_T446
GTF2F1_S156
IMPDH2_S416
LIG1_S51
LIG1_S66_S76
LIG1_T233
MPG_T67
NELFE_S51
NELFE_S115
NELFE_S251
NELFE_T91
NT5C_S182
RNMT_T77
RPA2_S174
SEC61A1_S408
SF3A3_T475
SRSF6_S316
SRSF6_S259
SRSF6_S280
SRSF6_S119
SSRP1_S673
SSRP1_S621
TP53_S315
UMPS_S214
USP11_S948_S953
ARL6IP1
POLR2I
TP53
CMPK2
NME3
NT5C3A
PDE4B
POLD4
RAD52
REV3L
SNAPC4
TAF13
TMED2
VPS37D
CETN2
LIG1
SRSF6
G
Protein Ubiquitination
hpi
3 6 12 24 48
AMFR_S509
BRCAT_S1497
HUWE1_S3662
HUWE1_T2751
HUWE1_T2747
HUWE1_T2738
HUWE1_S2887
HUWE1_S1368
PCNP_S77
SAE1_S185
UBE2V1_S146
UBE4B_S238
UBE4B_S803
BCL10_S138
GTPBP4
UBE2C
BCL10
GTPBP4
ANAPC11
CBL
PRKN
RNF144B
TRAF6
TRIM23
H
Regulation of Protein Mono and Polyubiquitination
hpi
3 6 12 24 48
DDX3X_S614
DDX3X_S594
FBXO4_T/S
OTUB1_S16
PPIA_S40
TRIP12_S85
TRIP12_S100
TRIP12_S127
TRIP12_S997
TRIP12_S1036
TRIP12_S1427
UBE2V1_S146
UBR5_S636
PARP10_S1011
PARP10
MARCHF7
NMI
NOD2
OTUB2
PRKN
PTPN22
FANCM
log2(fold change)
-0.5
0
0.5
Phosphoproteome
Proteome
Transcriptome
I
2
1
0
−1
−2
log2(fold change)
PJA2
RLIM
RNF14
RNF169
RNF5
UBE2J1
UBE2S
UBE2T
WWP2
3
6
12
24
48
hpi
J
hpi
3
6
12
24
48
Regulation of cell cycle process
PTTG1
UBE2C
TP53
CKS1B
MIS18BP1
FAM83D
CDK1
TIPIN
AHR
ENSA
UBE2S
AURKA
CKAP2
RACGAP1
NUSAP1
KIF2C
CCNB1
ECT2
CDC20
ASPM
DLGAP5
ANLN
TPX2
GMNN
PRC1
log2(fold change)
−1.5
0
1.5
